# CAR T cell foundation model predicts immunotherapy response

**DOI:** 10.64898/2026.07.11.737994

**Authors:** Yawei Li, Deyu Fang, Chengsheng Mao, Xin Wu, Yuan Luo

## Abstract

Single-cell transcriptomics resolves CAR T-cell states, yet translating heterogeneous cellular signals into patient-level therapeutic response remains challenging. Existing studies primarily identify response-associated genes or cell populations through experimental and statistical analyses, but few predictive frameworks integrate gene-level structure with clinical outcomes. Here, we present gANCHOR, a T-cell foundation model built on a hierarchical hypergraph attention framework combining biologically informed representation learning with patient-level response prediction. By encoding gene-pathway relationships, gANCHOR learns pathway-aware cell embeddings that improve biological conservation and batch robustness. A cell-to-patient attention module then aggregates cellular information to infer therapeutic response. Across benchmark datasets, gANCHOR achieved the strongest overall performance in biological conservation and batch-correction assessments. In response prediction across 161 patients from five CAR T-cell studies, gANCHOR achieved an F1 score of 0.87, outperforming benchmarked single-cell foundation models. gANCHOR also identified reproducible response- and non-response-associated gene programs, providing interpretable biological insights into CAR T-cell efficacy and resistance.

## Introduction

Chimeric antigen receptor (CAR) T-cell therapy, in which patient-derived T lymphocytes are genetically engineered to target tumor-associated antigens, has transformed cancer treatment by producing unprecedented responses in otherwise refractory hematological malignancies ^1^. By combining an antibody-derived antigen recognition domain with intracellular T-cell signaling modules, CAR constructs enable antigen-specific cytotoxicity independent of major histocompatibility complex restriction ^2^. Over the past decade, anti-CD19 CAR T therapies have achieved durable remissions in patients with refractory disease, leading to multiple regulatory approvals and widespread clinical adoption ^3-6^. Despite these successes, substantial variability in treatment outcomes, including relapses and treatment-associated toxicities, highlights the need for a deeper understanding of the biological determinants driving CAR T-cell therapeutic efficacy ^7^.

Advances in single-cell RNA sequencing (scRNA-seq) have provided high-resolution characterization of cellular heterogeneity in cancer and the tumor microenvironment ^8^. Unlike bulk profiling, which averages gene expression across mixed populations, scRNA-seq resolves transcriptomes at single-cell resolution, enabling the identification of rare cell subsets ^9^, lineage trajectories ^10^, and dynamic cell-cell interactions ^11^ that influence therapeutic response. In immuno-oncology, these technologies have revealed diverse T cell differentiation states, exhaustion programs, and functional heterogeneity within tumors, offering critical insights into mechanisms underlying response and resistance to therapies such as immune checkpoint blockade and CAR T-cell therapy ^12,13^. As scRNA-seq datasets continue to expand in scale and diversity, they represent a powerful foundation for developing predictive and mechanistic models of tumor-immune interactions.

Despite these advances, predictive modeling of CAR T-cell therapeutic response remains limited. Current clinical stratification strategies rely largely on bulk-derived biomarkers, which capture only a fraction of the variability in patient outcomes ^14-16^. Although recent scRNA-seq studies have identified transcriptional programs associated with CAR T-cell differentiation, persistence, and dysfunction ^17-30^, these analyses are predominantly descriptive and correlation-based. In parallel, recent single-cell foundation models have demonstrated that large-scale pretraining on diverse transcriptomic datasets can yield transferable cellular representations for downstream biological tasks. However, few, if any, deep learning frameworks directly leverage scRNA-seq data for patient-level prediction of CAR T-cell therapeutic outcomes. Bridging this gap enables personalized immunotherapy and uncovering molecular features that drive response and resistance.

Here, we present gANCHOR, a deep learning framework designed to predict CAR T-cell therapeutic outcomes from single-cell transcriptomic data while providing interpretable biological insights. gANCHOR employs a hypergraph-based neural architecture in which genes are connected through curated pathway relationships, enabling structured modeling of gene-gene dependencies within each cell. Pretrained on approximately 1.8 million T and CAR T cells across multiple datasets (**Table S1, S2**), the model learns biologically informed representations that capture T cell heterogeneity while mitigating batch effects. These embeddings are subsequently used for patient-level prediction, where gANCHOR outperforms state-of-the-art approaches. Importantly, the pretrained embeddings and model weights are transferable, enabling gANCHOR to be applied to new scRNA-seq datasets without retraining the full model.

Beyond CAR T-cell therapy, this design provides broader immunological studies where T-cell heterogeneity plays central roles, supporting biomarker discovery, patient stratification, and task-specific fine-tuning.

## Results

### gANCHOR establishes a hierarchical framework for CAR T response prediction

We developed gANCHOR (<u>g</u>raph <u>A</u>ttention <u>N</u>etwork for <u>C</u>AR-T <u>H</u>ierarchical <u>O</u>utcome <u>R</u>easoning), a deep learning framework designed to infer patient-level therapeutic outcomes from CAR T single-cell transcriptomes (**Figure 1**). gANCHOR adopts a hierarchical two-module architecture in which Module I learns biologically informed cell embeddings and Module II performs attention-based patient-level response prediction. This framework enables modeling of pathway-guided gene relationships within cells and cellular heterogeneity across patients while preserving interpretability through hierarchical attention mechanisms.

**Figure 1.**
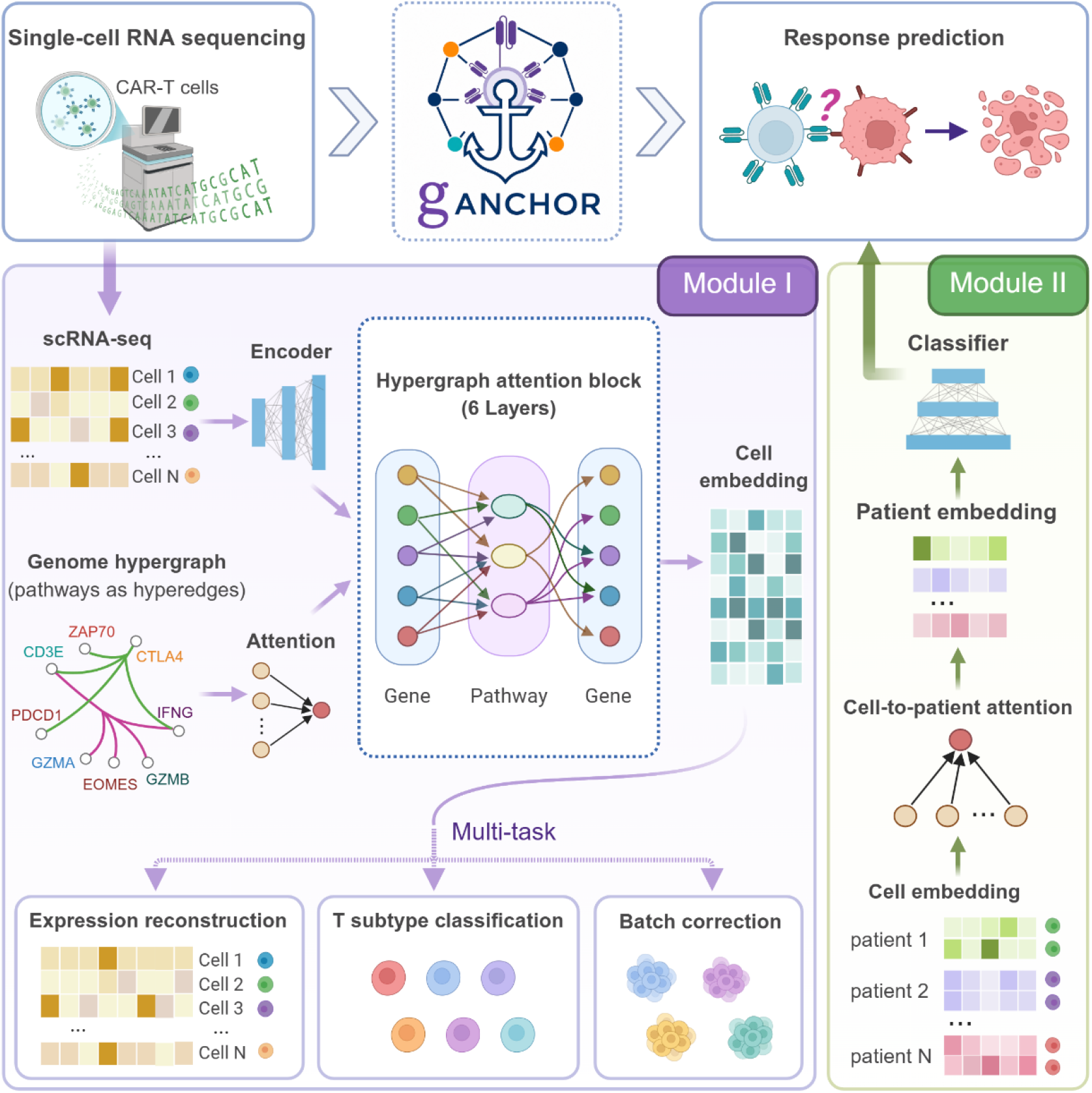
gANCHOR framework for hierarchical modeling of CAR-T single-cell data and patient response prediction. gANCHOR is a two-module architecture that integrates gene-pathway structure and cellular heterogeneity to predict patient-level therapeutic outcomes from single-cell RNA-seq data. Module I (foundation model) learns biologically informed cell embeddings by incorporating a curated gene-pathway hypergraph. A gene encoder performs attention-based message passing between genes and pathways to generate pathway-aware gene representations, which are combined with single-cell expression profiles and processed through stacked hypergraph attention blocks to model structured intra-cellular dependencies. A multi-task decoder reconstructs gene expression and predicts cell types, while adversarial training removes batch effects, yielding transferable, biologically meaningful cell embeddings. Module II aggregates cell embeddings via a cell-to-patient attention mechanism, assigning adaptive weights to individual cells and generating patient-level representations for response classification. Training is performed in two stages: Module I is pretrained with multi-task and batch-adversarial objectives, and the resulting embeddings are fixed for supervised training of Module II. This design enables interpretable modeling of gene-pathway interactions, cell-state heterogeneity, and their contributions to clinical response.

Module I serves as a single-cell foundation model trained on integrated CAR T-cell and reference T-cell datasets. Rather than treating genes as independent features, it incorporates curated gene-pathway relationships through a hypergraph attention framework. This architecture involves a gene encoder, a cell encoder, and a cell decoder. In the gene encoder, a bipartite gene-pathway graph is constructed using Molecular Signatures Database (MSigDB) ^31^, and hypergraph attention layers generate pathway-aware gene embeddings as structured biological priors. The cell encoder then integrates single-cell gene expression with gene embeddings to capture pathway-informed transcriptional dependencies while preserving cell-to-cell heterogeneity. The cell decoder further optimizes these representations through multi-task objectives, including gene expression reconstruction and cell-type prediction, while an adversarial batch discriminator reduces dataset-specific technical variation. This training strategy yields transferable and batch-corrected cell embeddings.

Module II uses the learned cell embeddings to predict patient-level response. A cell-to-patient attention mechanism dynamically weights individual cells and aggregates them into a unified patient representation for downstream classification. Training is performed in two independent stages. Module I is first pretrained to learn transferable cell embeddings, after which the encoder is fixed and Module II is then trained separately for patient-level prediction. This strategy stabilizes optimization and reduces overfitting to limited clinical outcome labels.

### gANCHOR achieves robust biological conservation and batch correction

High-quality cell representations are essential for both patient-level prediction as well as downstream biological interpretation. We therefore benchmarked gANCHOR against several batch-correction methods (Combat ^32^, MNN ^33^, scVI ^34^ and Scanorama ^35^) and single-cell foundation models (Geneformer ^36^, scGPT ^37^, scMulan ^38^, scFoundation ^39^ and GenePT ^40^). UMAP visualization revealed that, although batch effects were not completely eliminated, gANCHOR achieved improved integration between T-cell and CAR T-cell datasets while preserving biologically meaningful structure (**Figure 2A,B; Figure S2**). Within T-cell datasets, gANCHOR embeddings clustered predominantly by cell type rather than sequencing platform, suggesting effective batch mixing while maintaining cellular identity.

**Figure 2.**
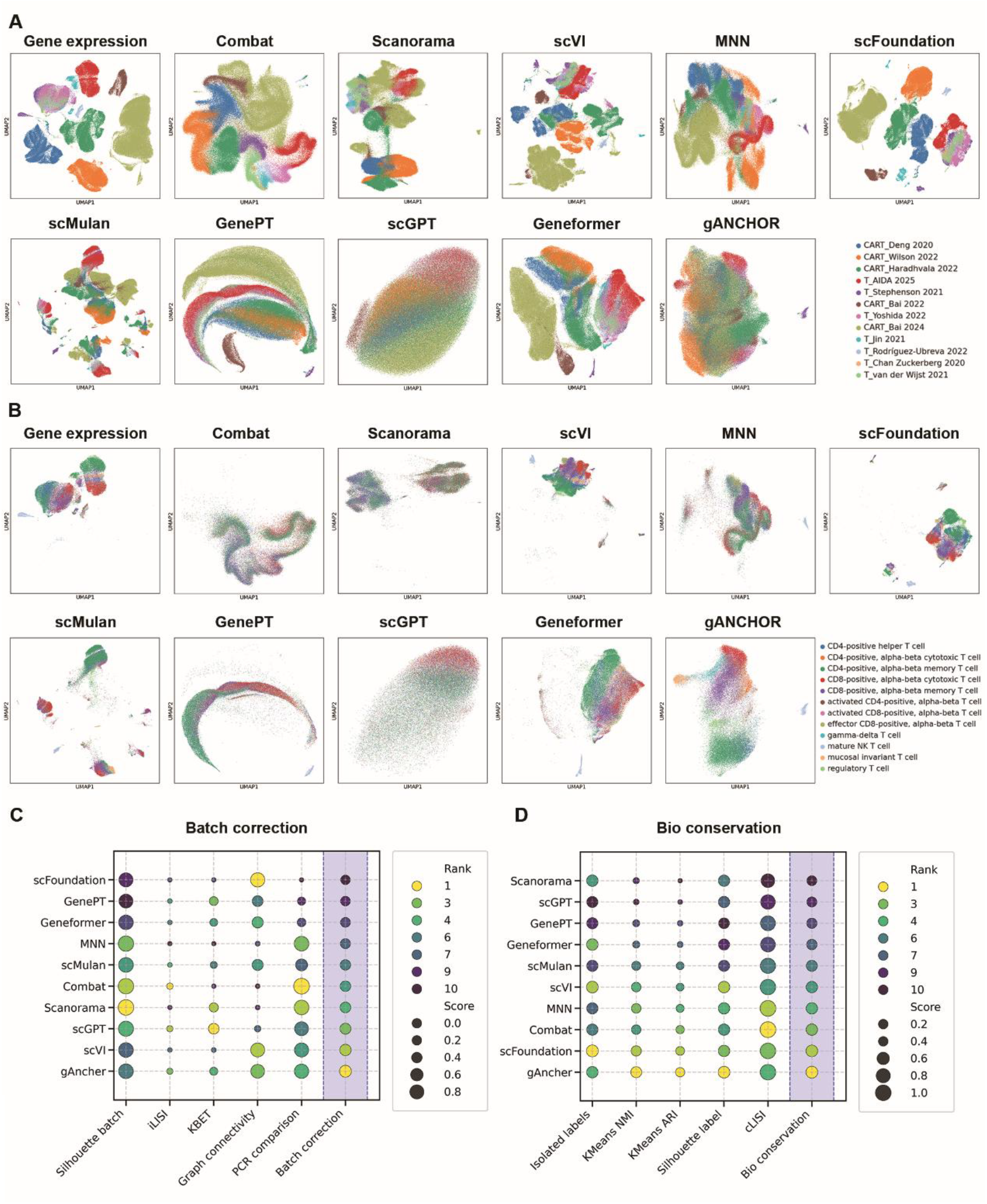
Benchmarking of cell embedding quality across models. (**A, B**) UMAP visualizations of cell embeddings generated from raw gene expression, Combat, Scanorama, scVI, MNN, scFoundation, scMulan, GenePT, scGPT, Geneformer, and gANCHOR. Cells are colored by dataset (**A**) and T cell subtype (**B**). (**C, D**) Quantitative evaluation of embedding performance across models using a comprehensive set of metrics. Batch correction metrics (**C**) include silhouette score for batch mixing, integration local inverse Simpson’s index (iLISI), k-nearest-neighbor batch effect test (kBET), graph connectivity, and principal component regression (PCR) comparison. Dot size represents the normalized metric score, and color indicates relative ranking across methods. Biological conservation metrics (**D**) include isolated label score, normalized mutual information (NMI), adjusted Rand index (ARI), silhouette score for cell types, and cell-type local inverse Simpson’s index (cLISI). gANCHOR achieves the highest overall performance, demonstrating strong and consistent results across both biological conservation and batch correction metrics.

To quantitatively evaluate embedding quality, we computed a comprehensive set of metrics assessing both biological conservation and batch correction performance ^41^. These involve isolated label score, ARI, NMI, ASW, LISI, graph connectivity, kBET, and PCR comparison. Across these metrics, gANCHOR demonstrated the strongest overall performance across both metric categories (**Figure 2C,D**). Specifically, gANCHOR ranked among the top-performing methods for biological conservation, with strong isolated label score, NMI, ARI, and cell-type silhouette metrics, indicating preserved cellular identity and subtype structure. It also achieved the highest aggregate batch-correction score, supporting effective integration across studies while preserving local biological structure.

Notably, several methods preferentially optimized one objective at the expense of the other. Conventional batch-correction approaches generally improved dataset mixing but often showed weaker preservation of biological structure, whereas some foundation models achieved strong biological conservation while exhibiting more limited batch correction. In contrast, gANCHOR maintained a balanced trade-off between these competing objectives and achieved the highest aggregate performance overall.

### gANCHOR improves patient-level CAR T response prediction

Given the strong biological conservation and batch-correction performance of gANCHOR embeddings, we next evaluated whether these representations improve patient-level CAR T therapy response prediction (**Figure 3A, B**). For all single cell foundation models, including gANCHOR, pretrained cell embeddings were used as fixed inputs to a unified prediction framework (Module II), ensuring that performance differences primarily reflect representation quality rather than classifier design. Raw gene expression profiles were also included as an additional baseline, allowing direct comparison between learned representations and unprocessed transcriptomic features.

**Figure 3.**
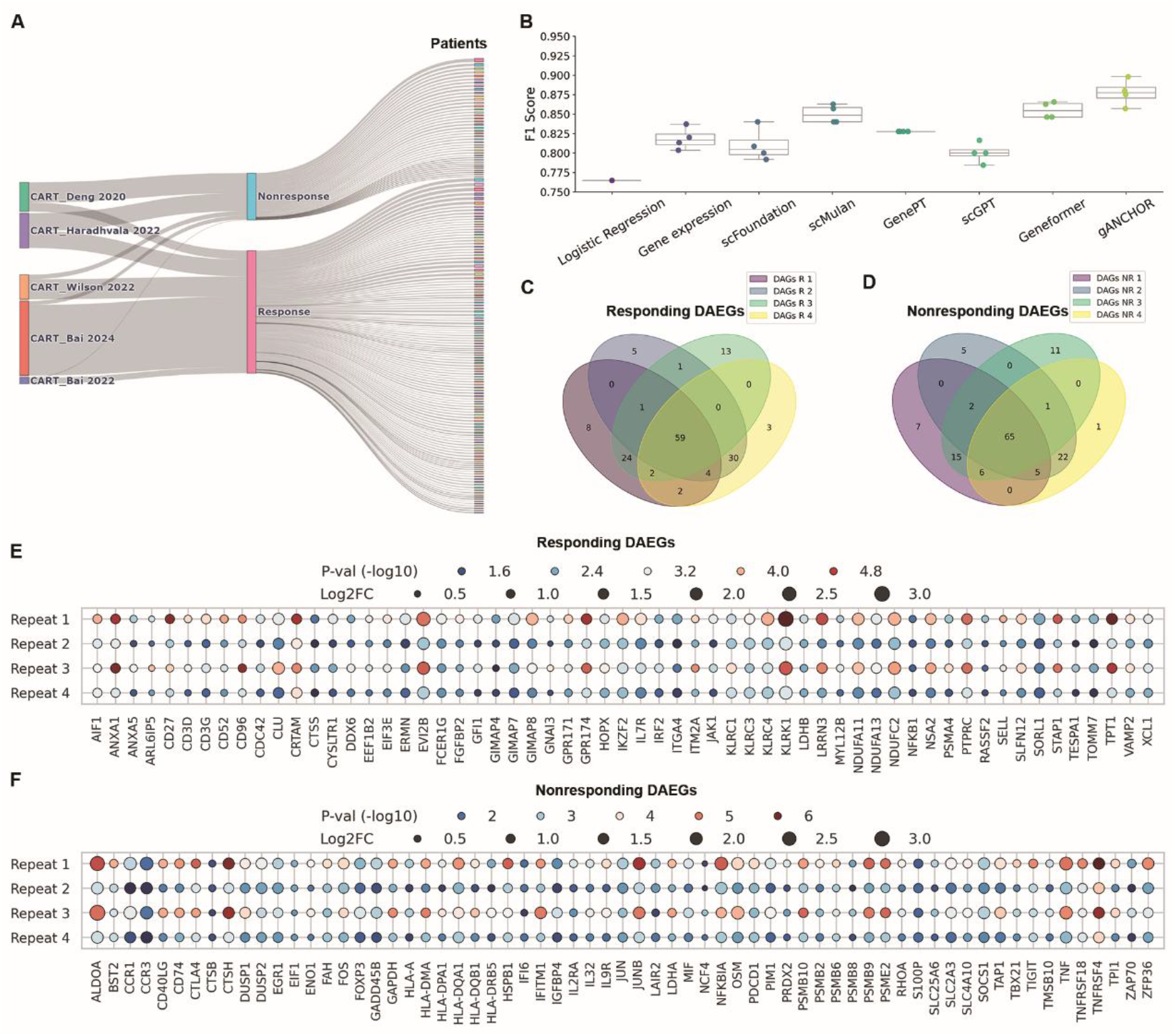
gANCHOR improves patient-level response prediction and reveals reproducible response-associated gene programs. (**A**) Sankey diagram illustrating the distribution of CAR T cells across datasets, clinical response, and individual patients. (**B**) Patient-level response prediction performance measured by F1 score. All models, except logistic regression, were evaluated across four independent runs with different random initializations. gANCHOR achieves the highest mean performance with low variability and shows a statistically significant improvement over all baseline models (permutation test, P < 0.05), indicating robust and reproducible predictive capability. (**C, D**) Venn diagrams showing the overlap of the top 100 differential attention-enriched genes (DAEGs) identified across four independent runs in the responding (**C**) and nonresponding (**D**) groups. A substantial fraction of genes is consistently recovered across all runs, including 59 genes in the responding group and 65 genes in the nonresponding group, demonstrating high reproducibility of the identified gene signatures. (**E, F**) DAEGs in the responding (**E**) and nonresponding (**F**) groups, restricted to genes consistently identified across all runs. Dot size represents log2 fold change (log2FC), reflecting the magnitude of increased gene contribution, and color indicates statistical significance (−log10 P value). These patterns highlight distinct and robust gene programs associated with favorable and unfavorable clinical outcomes.

Across all models and input settings, gANCHOR consistently achieved the best predictive performance on the held-out test data. It attained the highest F1 score (**Figure 3B; Figure S3**) while maintaining robust performance across complementary evaluation metrics, including precision, recall, AUROC, and AUPRC (**Figure S4**), demonstrating balanced discrimination between responder and non-responder patients. Notably, raw gene expression profiles performed substantially worse than embedding-based approaches, highlighting the importance of representation learning for capturing predictive cellular features. To further assess the contribution of cell-level modeling, we performed an additional ablation by collapsing single-cell data into patient-level features via mean aggregation of gene expression across all cells, followed by logistic regression for response prediction (**Figure 3B**). This simplified baseline achieved the lowest F1 score among all evaluated methods, indicating that naive aggregation fails to capture the cellular heterogeneity required for accurate response prediction.

To assess the statistical significance of these improvements, we performed a permutation test by randomly reassigning test F1 scores between models and recomputing the difference in mean performance. gANCHOR demonstrated a statistically significant advantage over all baseline models, indicating that the observed performance gains are unlikely to arise from stochastic variation across runs. Because all methods were evaluated using the same downstream prediction framework, these improvements can be primarily attributed to differences in representation quality . Together, these results demonstrate that gANCHOR provides a more discriminative and biologically informative representation space for CAR T therapy response prediction.

### gANCHOR identifies reproducible gene programs associated with clinical response

Having established that gANCHOR accurately predicts patient-level therapeutic response, we next investigated the molecular programs underlying these predictions by quantifying gene-level contributions through the hierarchical attention structure of the model. Patient-specific gene attention scores were computed by propagating importance from genes to cells and subsequently to patients, enabling identification of differential attention-enriched genes (DAEGs) between responding and non-responding patients (see Methods). Across four independent training repeats (**Table S3**), we identified top 100 DAEGs for each group, with 59 genes in the responding group (**Figure 3C, E**) and 65 genes in the non-responding group (**Figure 3D, F**) consistently identified across all runs. These gene sets reflect distinct immune states and provide a robust basis for interpreting the molecular determinants of CAR T-cell therapeutic outcomes.

In the responding group, DAEGs were enriched for genes associated with T cell activation, memory formation, and cytotoxic effector function (**Figure 3E**). For example, *SELL* (CD62L), *IL7R* (CD127), and *CD27* are canonical markers of central memory and stem-like T cells that have been consistently linked to improved CAR T expansion and persistence. *XCL1* is involved in chemokine-mediated immune cell recruitment and coordination, whereas *KLRK1* (NKG2D) and *FGFBP2* mark highly cytotoxic effector programs. *PTPRC* underscores the importance of intact T-cell receptor signaling in therapeutic efficacy. Together, these genes indicate a functionally competent, memory-enriched, and cytotoxic CAR T-cell state associated with favorable outcomes.

In contrast, DAEGs enriched in non-responding patients exhibited signatures of T cell dysfunction and exhaustion (**Figure 3F**). Immune checkpoint molecules such as *PDCD1* (PD-1), *CTLA4*, and *TIGIT* are well-established exhausted T cell markers, and have been linked to impaired CAR T expansion and reduced antitumor activity. Transcription factors *JUN, JUNB*, and *FOS*, components of the AP-1 complex, are associated with chronic activation and stress responses. Enrichment of *FOXP3* suggests increased regulatory T cell-like programs contributing to immunosuppression.

Importantly, these response-associated gene signatures are highly consistent with observations across independent CAR T scRNA-seq studies included in this work. Memory-associated genes identified in the responder group, including *SELL, IL7R*, and *CD27*, align with previously reported features of effective infusion products and durable response trajectories ^17,19^. Conversely, the enrichment of exhaustion-associated markers such as *PDCD1, CTLA4*, and *TIGIT* in the non-responder group recapitulates transcriptional programs linked to treatment failure and relapse ^18,29^. This concordance reflects a broader memory-versus-dysfunction axis, supported by regulatory and stress-associated genes such as *FOXP3, JUN*, and *FOS* ^30^. These findings demonstrate that gANCHOR recovers biologically meaningful response-associated programs while providing a quantitative framework for interpreting mechanisms associated with CAR T-cell efficacy and failure.

### gANCHOR learns biologically structured and interpretable gene and cell representations

Beyond patient-level response prediction, we further examined whether gANCHOR captures biologically meaningful cellular and gene-level organization within the learned representation space. To assess cell-level representations, we examined the model’s ability to classify T cell subtypes. The model achieved robust classification performance, with the confusion matrix showing accurate discrimination of major T cell populations while preserving expected similarities among closely related states (**Figure 4A, B; Figure S5**). These results indicate that the learned cell embeddings encode biologically relevant transcriptional programs and support reliable cell identity resolution.

**Figure 4.**
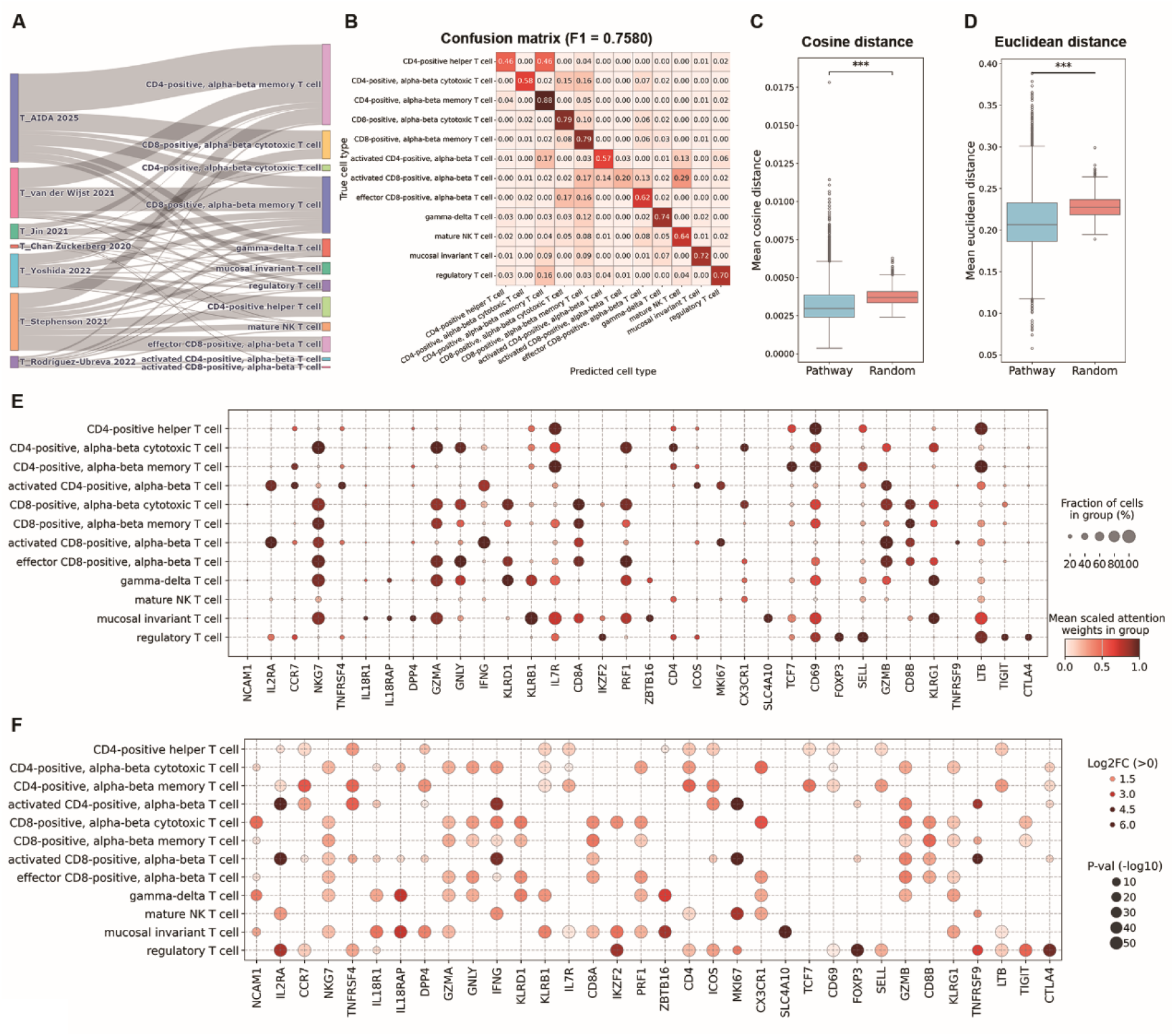
gANCHOR learns biologically informed gene and cell representations. (**A**) Sankey diagram illustrating the distribution of cells across dataset batches and annotated T cell types. (**B**) Confusion matrix showing the predictive performance of gANCHOR for T cell type classification in the test dataset (F1 = 0.7580). Values are normalized by the total number of cells in each ground truth class, enabling comparison of classification accuracy across cell types. (**C, D**) Mean pairwise cosine (**C**) and Euclidean (**D**) distances of gene embeddings for genes within the same pathways compared to randomly selected gene sets. Genes within the same pathway exhibit significantly smaller distances than those in randomly sampled sets (1,000 gene sets, Wilcoxon rank-sum test, *** P < 0.001), indicating that the learned gene embeddings preserve pathway-level organization and capture biologically meaningful relationships. (**E, F**) Differential attention-enriched genes of T cell subtype marker genes across subtypes, derived from the gene-to-cell attention mechanism. (**E**) Scaled attention weights and fraction of cells with non-zero attention (expression > 0), reflecting subtype-specific gene importance. (**F**) Log2 fold change (log2FC) and statistical significance (−log10 P-value); only genes with log2FC > 0 (increased attention) are shown. These patterns highlight distinct transcriptional programs associated with different T cell subtypes, with canonical marker genes consistently showing increased attention (log2FC > 0) in their corresponding cell types, supporting the biological fidelity of the learned representations.

To assess gene-level organization, we analyzed relationships among genes in the learned embedding space by comparing pairwise distances between genes belonging to the same MSigDB pathways with distances from randomly sampled gene sets. Genes within the same pathway exhibited significantly smaller pairwise distances (Wilcoxon rank-sum test, P < 0.001) (**Figure 4C, D; Figure S6**), indicating that the learned embeddings preserve pathway-level functional coherence. This structured organization suggests that genes participating in shared biological processes are embedded in close proximity within the latent space.

To further evaluate interpretability, we analyzed gene-to-cell attention weights and identified DAEGs for each T cell subtype (see Methods). These subtype-specific DAEGs revealed strong concordance with established marker genes, with most canonical markers exhibiting positive log2 fold change and high statistical significance, indicating consistently elevated attention within their corresponding cell types (**Figure 4E, F**). Cytotoxic T cell subsets, including CD8-positive and CD4-positive cytotoxic populations, showed increased attention to effector molecules such as *GZMB, PRF1, GNLY*, and *NKG7*, consistent with cytolytic function. Memory T cell subsets displayed enriched attention to *IL7R* and *CCR7*, reflecting lymphoid homing and long-term persistence. Regulatory T cells exhibited strong and specific attention to *FOXP3, IL2RA*, and *CTLA4*, capturing their immunosuppressive phenotype. Activated T cell states were characterized by elevated attention to activation markers such as *CD69* and *IL2RA*, while innate-like T cell populations, including mucosal invariant T cells and natural killer T cells, show increased attention to *KLRB1* and *ZBTB16*.

To determine whether these biologically structured representations depend on the curated pathway priors rather than the hypergraph attention architecture alone, we performed an ablation analysis in which gene-to-pathway assignments were randomly shuffled while preserving the overall hypergraph topology. Compared with the original MSigDB-derived pathway graph, the shuffled model exhibited reduced biological conservation and batch-correction performance (**Figure S7A-D**), indicating that disrupting biologically meaningful gene-pathway relationships compromises the quality of the learned representations. Consistent with this reduction in representation quality, the shuffled model also assigned weaker attention to canonical subtype marker genes, reflected by lower attention-associated log2 fold changes across T-cell populations (**Figure S7E, F**). Together, these findings demonstrate that the biological interpretability and representational quality of gANCHOR depend not only on the hierarchical attention architecture but also on biologically meaningful gene–pathway organization encoded by curated pathway annotations.

### Attention-derived representations preserve biologically structured cellular states

To assess whether the learned attention mechanisms provide informative representations beyond embeddings, we evaluated gene-to-cell attention outputs as an alternative feature space for characterizing cellular states. For each cell, attention-weighted gene representations were extracted from the gene-to-cell attention module and used for downstream analyses. These representations revealed coherent organization of annotated T-cell subtypes while improving dataset mixing (**Figure S8A-C**). In the integrated reference T-cell and CAR T-cell dataset, attention-based representations preserved T-cell structure while retaining CAR T-cell heterogeneity, suggesting that the model avoids overcorrection under domain shift.

Quantitative evaluation further showed that attention-derived representations achieved biological conservation comparable to normalized gene expression while substantially improving batch correction (**Figure S8D**). Although learned cell embeddings remained the best-performing representation overall, attention-derived features consistently outperformed raw gene expression in overall integration quality, suggesting that the gene-to-cell attention mechanism preserves biologically meaningful cellular structure while reducing technical variation across datasets and retaining gene-level resolution.

We next investigated whether attention-derived representations provide biological information beyond conventional differential expression analysis. Using identical statistical procedures, we compared DAEGs with differential expression genes (DEGs) identified from gene expression. At the cell-type level, where batch effects are relatively modest (**Figure 2A**), both approaches recovered canonical subtype marker genes with comparable accuracy (**Figure 3E,F** and **Figure S9**), indicating that attention faithfully preserves established transcriptional programs while remaining consistent with conventional expression-based analysis. In contrast, for patient response analysis, where CAR T-cell datasets exhibit substantially stronger batch effects (**Figure 2A**), the two approaches diverged considerably (**Table S4**). Conventional DEGs were dominated by broadly expressed housekeeping, metabolic, and antigen-presentation genes, whereas DAEGs preferentially highlighted genes with well-established roles in CAR T-cell biology, including cytotoxicity, immune regulation, and T-cell exhaustion. Notably, several of these biologically relevant genes were absent from the top-ranked DEGs despite being consistently prioritized by the attention model. These findings suggest that attention-derived representations suppress technical and dataset-specific variation while amplifying biologically informative transcriptional programs, providing a more interpretable and clinically relevant feature space than conventional differential expression analysis.

## Discussion

gANCHOR establishes a hierarchical hypergraph attention framework that connects single-cell transcriptomic representation learning with patient-level therapeutic outcome prediction in CAR T-cell therapy. By encoding curated gene-pathway relationships as a bipartite hypergraph, the foundation module learns pathway-aware gene and cell embeddings that preserve coordinated functional programs while reducing batch effects. The cell-to-patient attention module then aggregates heterogeneous cellular states into patient-level embeddings for accurate and interpretable response prediction. Under a unified downstream architecture, gANCHOR consistently achieves improved predictive performance. The hierarchical attention structure propagates feature importance from genes to cells and patients, allowing identification of gene programs associated with therapeutic response. The resulting gene signatures show concordance with reported CAR T-cell functional states, including activation, memory, cytotoxicity, and exhaustion, indicating that gANCHOR captures biologically meaningful signals relevant to therapeutic outcomes.

Beyond response prediction, gANCHOR provides a generalizable framework for linking single-cell molecular representations to clinically relevant phenotypes across heterogeneous T-cell and CAR T-cell scRNA-seq datasets. By integrating pathway-guided representation learning with hierarchical attention, the framework enables interpretable mapping from molecular features to cellular states and patient-level predictions while maintaining robustness across diverse studies and sequencing platforms. This design supports both response prediction and identification of clinically relevant genes and pathways. As a T cell-focused foundation model, gANCHOR learns reusable representations from large-scale integrated T-cell and CAR T-cell transcriptomic datasets. These transferable representations and attention-derived scores provide useful features for biomarker discovery, patient stratification, and task-specific fine-tuning in related immunological settings where T-cell heterogeneity is central. More broadly, this study illustrates the feasibility of integrating structured biological priors with deep learning to bridge the gap between high-dimensional single-cell data and clinically actionable insights.

gANCHOR also has several limitations. The gene-pathway hypergraph relies on existing pathway annotations, which may be incomplete or biased toward well-characterized biological processes, potentially limiting the discovery of unannotated functional relationships. Although attention mechanisms improve interpretability, attention weights do not constitute direct evidence of causal regulation and require complementary experimental validation. In addition, the foundation and response modules are trained in separate stages; future work could explore end-to-end or task-adaptive fine-tuning strategies to better align representation learning with clinical prediction objectives. Extending gANCHOR to incorporate multi-omic data, spatial transcriptomics, and longitudinal sampling may further improve its ability to capture dynamic immune processes. Finally, validation in larger and more diverse patient cohorts will be necessary to assess generalizability and potential clinical utility.

## Methods

### T cell and CAR T cell single cell data collection and initial filtering

scRNA-seq datasets of T cells and CAR T cells were collected from twelve published studies spanning multiple disease contexts. All CAR T-cell samples used in this study were derived from pre-infusion material collected prior to clinical outcome assessment, ensuring that both representation pretraining (Module I) and patient-level response prediction (Module II) were performed exclusively using available transcriptomic information. These CAR T-cell datasets enabled characterization of transcriptional programs associated with subsequent therapeutic response ^17-19,29,30^. Reference T-cell datasets were obtained from large-scale atlases and disease-focused cohorts and were subsetted to retain annotated T-cell populations across diverse functional subtypes ^42-48^ through the CZ CELLxGENE database ^49^. All datasets were processed using a unified preprocessing workflow, including normalization to account for sequencing depth and harmonization to a shared gene space where necessary. For datasets lacking raw counts, expression values were transformed to ensure compatibility with downstream analysis. For further details on how each dataset was collected, curated, and processed, please see the Supplementary Information.

### Harmonization of patient-level clinical response labels

A total of 161 patients from five independent studies ^17-19,29,30^ were included for patient-level response prediction. Clinical outcome annotations from each study were harmonized into a unified binary classification framework consisting of response and nonresponse labels. Intermediate or study-specific response categories were systematically mapped to preserve consistency across cohorts while maintaining clinically meaningful distinctions. In total, 120 patients were classified as response and 41 as nonresponse (**Figure 3A, Table S1**).

For the Wilson et al. 2022 ^19^ cohort, 12 patients were retained after excluding two patients lacking pre-treatment scRNA-seq data and two lacking response annotations; 9 were classified as responders and 3 as nonresponders. In Haradhvala et al. 2022 ^18^, 31 of 32 patients were included after excluding one without scRNA-seq data. Patients labeled as response (n = 16) were assigned to the responder group, while those annotated as nonresponse (n = 14) or partial response (PR, n = 1) were grouped into nonresponder group. In Deng et al. 2020 ^17^, complete response (CR, n = 9) was mapped to response, whereas partial response (PR, n = 1), progressive disease (PD, n = 13), and death (n = 1) were categorized as nonresponse. The Bai et al. 2022 ^29^ cohort included 12 patients (10 response, 2 nonresponse) as originally reported. In Bai et al. 2024 ^30^, patients annotated as complete response (CR) or relapse after response (RL) were grouped as response (n = 76), whereas nonresponse (NR, n = 6) was assigned to the nonresponder group.

This harmonization strategy ensures a unified clinical endpoint across heterogeneous studies while preserving biologically and clinically relevant distinctions for downstream predictive modeling.

### Gene selection and pathway filtering

gANCHOR pretraining was based on genes with curated pathway annotations. We utilized the Molecular Signatures Database (MSigDB) ^31^ curated human gene set collection, specifically the C2 category, which comprises both Canonical Pathways (CP) and Chemical and Genetic Perturbations (CGP). These curated gene sets provide structured prior knowledge of gene-pathway relationships derived from experimentally validated biological processes and perturbation studies. The MSigDB C2 collection contains 21,587 unique genes. Genes lacking functional pathway annotations were excluded from downstream analysis. To ensure consistency across datasets, only genes that were detected across all pretraining T cell and CAR T-cell datasets were retained. Ribosomal genes were further removed to reduce potential confounding from technical variability and highly abundant housekeeping signals. After filtering, the final feature space consisted of 11,140 genes, which were used as for hypergraph construction and downstream representation learning.

### Pathway graph ablation analysis

To evaluate the contribution of curated biological pathway information, we performed an ablation experiment in which gene membership within MSigDB pathways was randomly shuffled while preserving the number of pathways and the number of genes assigned to each pathway. Module I was retrained using the shuffled gene-pathway incidence matrix under the same architecture and training procedure as the original model. Differential attention-enriched genes were subsequently identified using the same analysis pipeline. Biological interpretability was assessed by comparing the recovery of canonical subtype marker genes between the original MSigDB graph and the shuffled-pathway graph.

### Single-cell data normalization

scRNA-seq data preprocessing was performed using Scanpy. No additional cell- or gene-level filtering beyond the original study-specific quality-control procedures was applied beyond the criteria used in the original studies from which the datasets were obtained. Raw count matrices were first normalized to 10,000 total UMI counts per cell using scanpy.pp.normalize_total, followed by logarithmic transformation with scanpy.pp.log1p. To further mitigate gene-specific expression bias and technical variability associated with capture efficiency and sequencing depth, an additional gene-wise normalization step was applied. For each detected gene, the non-zero median expression value was calculated across all cells in the training datasets containing both T cells and CAR T cells. Gene expression values in each cell were then divided by the corresponding non-zero median. This procedure places genes with systematically different baseline expression levels onto a comparable scale while preserving biologically meaningful variation across cells, thereby providing stable inputs for downstream hypergraph-based representation learning.

### Harmonization of T-cell subtype annotations

All reference T cells used in this study were obtained from CZ CELLxGENE, where cell type annotations are based on the Cell Ontology ^50^. Because Cell Ontology annotations are hierarchical, individual cells may be assigned to highly specific subtypes nested within broader parent categories. To ensure consistency across datasets and compatibility with supervised classification tasks requiring mutually exclusive labels, we constructed a unified annotation framework consisting of 12 T-cell subtypes: CD4-positive helper T cell; CD4-positive, alpha-beta cytotoxic T cell; CD4-positive, alpha-beta memory T cell; CD8-positive, alpha-beta cytotoxic T cell; CD8-positive, alpha-beta memory T cell; activated CD4-positive, alpha-beta T cell; activated CD8-positive, alpha-beta T cell; effector CD8-positive, alpha-beta T cell; gamma-delta T cell; mature NK T cell; mucosal invariant T cell; and regulatory T cell (**Table S2**).

Fine-grained Cell Ontology annotations were systematically mapped to these unified categories. Specifically, central memory and effector memory CD4-positive T cells were consolidated into CD4-positive, alpha-beta memory T cells, and analogous CD8-positive memory subtypes, including terminally differentiated effector memory cells, were grouped into CD8-positive, alpha-beta memory T cells. Activated T cell annotations labeled with species-specific suffixes (e.g., “human”) were unified into their corresponding activated CD4-positive or CD8-positive categories. In addition, multiple regulatory T cell subtypes, including natural, double-negative, and naive regulatory T cells, were merged into a single regulatory T cell category. This harmonization strategy preserves major functional distinctions among T-cell states while ensuring annotation consistency across studies.

### Train-test data splitting strategy

To ensure rigorous and unbiased evaluation, data splitting was performed at both the cell and patient levels depending on the dataset type and modeling objective. For Module I pretraining, which integrates both reference T-cell and CAR T-cell datasets, distinct strategies were applied. In the reference T-cell datasets, 50,000 cells were randomly selected and held out as a test set, and all remaining cells were used for model training. For CAR T-cell datasets, splitting was performed at the patient level to prevent information leakage between samples derived from the same individual. Approximately 20% of patients (34 out of 161) were randomly selected as the test cohort, and all cells derived from these patients were excluded from training.

To maintain consistency between Module I and Module II, CAR T cells from the held-out test patients were excluded from all training procedures and used exclusively for final evaluation of patient-level response prediction. The same train, validation, and test splits were applied across gANCHOR and all benchmarked baseline models to ensure fair comparison of representation quality and predictive performance.

### Benchmark models and evaluation framework

We benchmarked gANCHOR against against four widely used batch-correction methods— Combat ^32^, MNN ^33^, scVI ^34^ and Scanorama ^35^—as well as five single-cell foundation models: Geneformer ^36^, scGPT ^37^, scMulan ^38^, scFoundation ^39^ and GenePT ^40^. Publicly available pretrained models and default parameter settings were used for all methods. The benchmarked foundation models included transformer-based cell embedding models (Geneformer, scGPT, scMulan, and scFoundation) as well as a gene embedding framework based on large-scale language models (GenePT). For transformer-based models, pretrained representations were directly extracted from gene expression profiles and used for downstream patient-level response prediction. For GenePT, pretrained gene embeddings were combined with expression profiles to generate cell representations (GenePT-w). Geneformer, scGPT, and scFoundation were fine-tuned according to their published protocols, whereas GenePT and scMulan were evaluated using their publicly available pretrained models. For scMulan, no additional training was performed because the model is designed for zero-shot downstream applications and the authors have not released a complete pretraining pipeline for continued model adaptation.

All models were evaluated using a unified downstream prediction framework, with pretrained embeddings treated as fixed inputs. Hyperparameter optimization was performed over learning rate and weight decay, and each parameter configuration was evaluated across four independent runs with different random seeds. The optimal hyperparameters were selected based on the highest mean binary F1 score on the validation set and subsequently evaluated on the held-out test cohort (**Figure S3**).

To systematically dissect the contribution of representation learning and cellular resolution, two complementary ablation settings were included. First, normalized gene expression profiles were directly used as inputs to the downstream prediction framework without representation learning. This setting provides a baseline to assess whether performance gains arise from representation learning rather than downstream modeling. Second, gene expression values were averaged across all cells within each patient to generate bulk-like patient-level profiles, which were then evaluated using logistic regression. This setting tests whether explicitly modeling cell-level variability provides additional predictive power beyond bulk-like representations. These ablations enabled assessment of the relative contributions of learned representations and explicit modeling of intra-patient cellular heterogeneity. Additional implementation details and parameter settings are provided in the Supplementary Information.

### Logistic regression baseline for patient-level prediction

For the aggregation-based baseline, gene expression values were averaged across all cells within each patient to generate patient-level gene expression profiles. The resulting patient-by-gene matrix was partitioned into training and test sets using the same split applied in the main benchmarking framework. Features were standardized using z-score normalization fitted on the training data and applied to the test data.

A logistic regression classifier was trained with hyperparameters optimized through grid search using 5-fold cross-validation on the training set. The search space included regularization strength (0.05, 0.1, 0.5, 1, 5, or 10), penalty type (L1 or L2), compatible solvers (liblinear, lbfgs, newton-cg, sag, or saga), and class weighting (none or balanced). The optimal configuration was selected based on cross-validation performance and evaluated on the held-out test set using F1 score. This baseline provides a reference framework for patient-level prediction without explicit modeling of single-cell heterogeneity or learned feature representations.

### Benchmark metrics and evaluation strategy on cell embeddings

To comprehensively evaluate the quality of learned cell embeddings, we adopted a benchmarking framework that jointly assesses biological conservation and batch correction, following established single-cell integration benchmarks ^41^. All metrics were computed either directly on the embedding space or on kNN graphs derived from the embedding. Biological conservation metrics evaluate whether integrated embeddings preserve cell identity structure, whereas batch correction metrics assess the extent to which cells from different datasets are appropriately mixed. All evaluations were performed using the same set of cells and annotations, including cell-type labels and batch (dataset) labels. Because CAR T cells lack unified subtype annotations across studies, biological conservation metrics were evaluated only on reference T-cell datasets, whereas batch correction metrics were computed on the combined T-cell and CAR T-cell datasets.

For biological conservation, we used clustering- and distance-based metrics, including normalized mutual information (NMI), adjusted Rand index (ARI), isolated label score, and average silhouette width (ASW). NMI and ARI quantify the agreement between unsupervised clustering (Louvain clustering on the embedding space) and ground-truth cell-type labels, where values closer to 1 indicate better recovery of cell identity structure. The isolated label score evaluates preservation of rare or underrepresented cell populations. ASW measures the separation of cell types based on intra-cluster and inter-cluster distances. For a given cell *i*, the silhouette width is defined as:

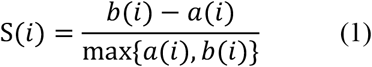

where *a*(*i*) is the average distance between cell *i* and cells of the same type, and *b*(*i*) is the minimum average distance to cells of different types. The ASW is obtained by averaging *s*(*i*) across all cells and is rescaling values to [0, 1], where higher scores indicate improved separation of biological cell types.

To evaluate batch correction, we used complementary metrics capturing both local neighborhood mixing and global batch structure. The batch ASW quantifies batch mixing by computing silhouette widths using batch labels instead of cell-type labels. To ensure t interpretability, silhouette values were inverted so that higher values correspond to improved batch mixing:

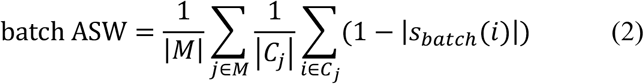

where *C*_*j*_ denotes cells of type *j*, and *M* is the set of all cell types. Values approaching 1 indicate effective removal of batch effects.

We further quantified neighborhood-level mixing using the local inverse Simpson’s Index (LISI), which measures label diversity within local neighborhoods in the kNN graph. Two variants were used: integration LISI (iLISI), which evaluates batch mixing and increases with improved integration, and cell-type LISI (cLISI), which evaluates preservation of biological structure and decreases when distinct cell identities remain well separated. LISI scores were computed from neighborhood graphs and rescaled to [0, 1] for comparability across datasets.

The k-nearest-neighbor batch effect test (kBET) was used to statistically assess whether local neighborhoods reflect the global batch composition. For each cell, kBET compares the batch distribution within its kNN neighborhood to the expected global distribution using a hypothesis test. The rejection rate across tested cells is used as the metric, and the final score is defined as 1 − rejection rate, such that higher values indicate improved batch mixing.

To evaluate preservation of cellular topology, we used the graph connectivity, which assesses whether cells of the same identity remain connected in the kNN graph. For each cell type *c*, the proportion of cells in the largest connected component was computed and averaged across all cell types:

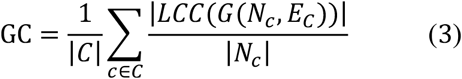

where LCC denotes the largest connected component. Higher values indicate improved preservation of within-cell-type connectivity structure.

Finally, principal component regression (PCR) was used to quantify residual batch-associated variation within the embedding space. Specifically, the variance explained by batch labels across principal components is computed as:

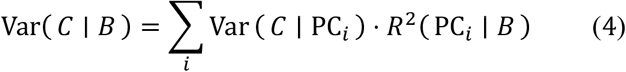

where *R*^2^(PC_*i*_ | *B*) quantifies the association between batch labels and each principal component. Lower values indicate reduced batch-driven variation and improved integration.

For each metric, scores were normalized and aggregated into two categories: biological conservation and batch correction. An overall performance score was then computed as the weighted average of these two categories, providing a balanced assessment of embedding quality across models.

### Assessment of pathway coherence in gene embedding space

To assess whether learned gene embeddings preserve biologically meaningful functional structure, we quantified pairwise distances among genes belonging to the same annotated pathways and compared these distances to randomly sampled gene sets. Pathway annotations were obtained from the MSigDB. For each pathway, mean pairwise distances among all pathway member genes were computed in the learned embedding space using both cosine and Euclidean distance metrics. As a background distribution, 1,000 random gene sets were generated by uniformly sampling genes from the complete gene list, with each random set matched in size to the corresponding pathway. Mean pairwise distances were computed for each random set using the same procedure. Distributions of pathway-based and random distances were then compared using the Wilcoxon rank-sum test to assess whether genes participating in shared biological pathways exhibit significantly closer organization in the embedding space than expected by chance.

### Benchmark metrics and evaluation strategy for patient-level response prediction

To assess binary CAR T therapy response prediction, we used the binary F1 score as the primary evaluation metric. The F1 score balances precision and recall and is particularly suitable for imbalanced classification settings in which responder and non-responder groups are unevenly distributed. Let true positives (TP), false positives (FP), false negatives (FN), and true negatives (TN) denote predicted response outcomes relative to ground truth labels. Precision and recall are defined as:

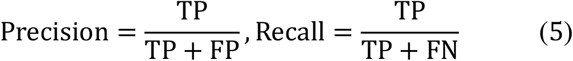

The binary F1 score is computed as the harmonic mean of precision and recall:

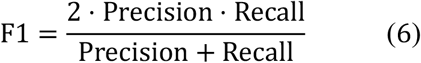

The F1 score ranges from 0 to 1, with higher values indicate improved discrimination between responder and non-responder groups.

Model performance was evaluated on a held-out test dataset using the optimal hyperparameter configuration selected from validation data (see Implementation and training details of Module II). To account for stochastic variability arising from random initialization, each model was evaluated across four independent runs with different random seeds. Test performance was summarized as the distribution of F1 scores across these runs, allowing assessment of both predictive accuracy and robustness.

For pairwise model comparisons, we performed a non-parametric permutation test by pooling test F1 scores from two models and randomly reassigning them into two groups matching the original group sizes. The difference in mean F1 score between the two groups was recomputed over 10,000 permutations to generate a null distribution. The empirical P-value was calculated as the proportion of permutations in which the permuted difference in means was greater than or equal to the observed difference. Statistical significance was defined as *P* < 0.05.

This evaluation framework enables robust comparison of patient-level predictive performance while avoiding strong distributional assumptions and accommodating the limited number of independent training runs.

### Computation and statistical analysis of differential attention-enriched genes

To quantify gene-level contributions across multiple biological resolutions, we leveraged the hierarchical attention architecture of gANCHOR to propagate feature importance from genes to cells and subsequently to higher-order structures, including cell types and patients.

We first extracted the gene-to-cell attention matrix from Module I, denoted as 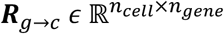, which captures the relative contribution of each gene within individual cells.

### Cell-type-level differential attention analysis

To characterize subtype-specific transcriptional programs, we analyzed gene attention patterns across annotated T-cell populations. For each cell, gene-level attention scores were obtained directly from ***R***_*g*→*c*_ . Cells were grouped according to their harmonized T-cell subtype annotations, and gene attention scores were aggregated within each subtype to generate subtype-specific gene importance profiles.

Differential attention-enriched genes at the cell-type level were identified using a one-versus-rest comparison framework. For each gene, a Mann-Whitney U test was performed to compare attention scores in cells belonging to a given subtype against all remaining cells.

Resulting P-values were adjusted for multiple testing using the Benjamini-Hochberg procedure to control the false discovery rate (FDR). Genes with significantly elevated attention weights in a given subtype were defined as subtype-associated DAEGs.

### Patient-level differential attention analysis

To connect gene-level signals to clinical outcomes, gene contributions was further propagated from cells to patients using the cell-to-patient attention matrix from Module II, denoted as 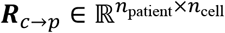, which represents the contribution of individual cells to patient-level predictions. Patient-specific gene contribution scores were computed as:

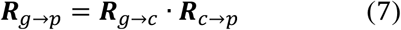

where 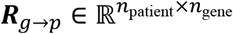 quantifies the contribution of each gene to the predicted therapeutic response of each patient.

To avoid information leakage from model evaluation, all differential analyses were restricted to patients in the training cohort. Patients were stratified into responding and nonresponding groups based on their clinical labels. To identify genes associated with differential clinical response, we applied a Mann-Whitney U test to compare contribution scores between the two groups, followed by Benjamini–Hochberg correction for multiple testing. Genes with significantly elevated contribution scores in responders were defined as responder-associated DAEGs, whereas those enriched in nonresponders were defined as nonresponder-associated DAEGs.

Genes were ranked based on their average contribution scores within each group, and the top 100 genes per group were selected for downstream analysis. To ensure robustness, this procedure was repeated across four independent training runs with different random seeds. Only genes consistently identified among the top 100 across all runs were retained as high-confidence DAEGs. Using this criterion, 59 responder-associated and 65 non-responder-associated DAEGs were identified.

### Differential expression analysis

For comparison with differential attention analysis, differential expression analysis was performed using Scanpy (scanpy.tl.rank_genes_groups) with the Wilcoxon rank-sum test. Cell-type-specific DEGs were identified using a one-versus-rest comparison across the 12 harmonized T-cell subtypes. For patient response analysis, only training-set CAR T cells were included to avoid information leakage. Cells were grouped according to the clinical response label of their corresponding patients, and DEGs between responders and nonresponders were identified using the same Wilcoxon framework. Genes were ranked according to the Scanpy ranking statistic, and the top-ranked genes were compared with canonical marker genes and literature-supported response-associated genes using the same evaluation procedure applied to DAEGs.

### Overall architecture of gANCHOR

gANCHOR consists of two sequential modules. Module I biologically informed cell representations from T-cell and CAR T-cell single-cell transcriptomes using a hierarchical hypergraph attention framework, whereas Module II aggregates cell embeddings into patient-level representations and therapeutic response prediction. The complete workflow is illustrated in **Figure 1**.

## Module I: Hypergraph-guided cell representation pretraining

### Input data representation

Let 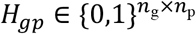 denote the gene–pathway incidence matrix, where *H*_*gp*_ = 1 indicates that gene *g* belongs to pathway *p*. Single-cell gene expression profiles are represented by the normalized expression matrix 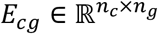, where *n*_*c*_ and *n*_*g*_ denote the numbers of cells and genes, respectively. Gene embeddings were initialized using a trainable embedding layer, producing 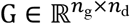. Cell-specific gene embeddings were constructed by broadcasting gene embeddings across cells and combining them with normalized expression values through element-wise multiplication, yielding 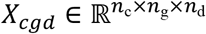, which serves as the input to the hierarchical graph-attention encoder. Following our notation convention, subscripts denote tensor dimensions; for example, *X*_*cgd*_ denotes a tensor of shape [*n*_*c*_, *n*_*g*_, *n*_*d*_]. Module I aims to learn pathway-aware gene and cell embeddings by integrating curated pathway structure with single-cell transcriptional profiles.

### Initial gene embedding via hypergraph graph attention

We constructed a gene-pathway bipartite hypergraph in which genes are represented as nodes and biological pathways are modeled as hyperedges connecting their member genes according to the incidence matrix *H*_*gp*_. To learn pathway-aware gene representations, we adopt a strongly dual-attention hypergraph message passing strategy inspired by SHINE ^51^. This framework allows genes to exchange information through shared pathway membership, capturing coordinated biological functions.

At each hypergraph attention layer, information is propagated bidirectionally between genes and pathways. First, for each pathway *p*, attention over connected genes quantifies the relative contribution of each gene to the functional identity of that pathway. These gene-to-pathway attention weights are normalized across member genes and used to update pathway embeddings by weighted aggregation of connected gene features.

Next, for each gene *g*, attention over connected pathways quantifies the relative contribution of each pathway to the functional context of that gene. These pathway-to-gene attention weights are normalized across connected pathways and used to update gene embeddings by weighted aggregation of pathway representations. Biologically, this step allows each gene to integrate contextual information from multiple biological pathways in which they participate.

After two stacked dual-attention hypergraph layers, the model outputs final structural gene embeddings 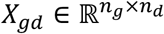, which encode global functional relationships among genes induced by shared pathway membership. These embeddings serve as biologically informed priors for subsequent cell-level modeling.

### Expression value encoding and fusion

Normalized expression values *E*_*cg*_ were transformed by a multilayer perceptron (MLP) to generate expression embeddings:

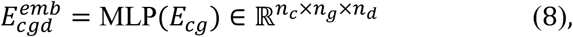

The pathway-derived gene embedding *G*_*gd*_ were broadcast across cells and combined with expression embeddings through element-wise multiplication:

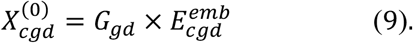

### Hierarchical hypergraph attention encoder

Within each hypergraph attention block, we perform a two-stage hierarchical attention procedure consisting of gene-to-pathway aggregation followed by pathway-to-gene redistribution. This design explicitly incorporates pathway structure into representation learning, enabling the model to capture coordinated regulation of genes participating in shared biological programs.

#### 1. Pathway attention (gene → pathway message passing)

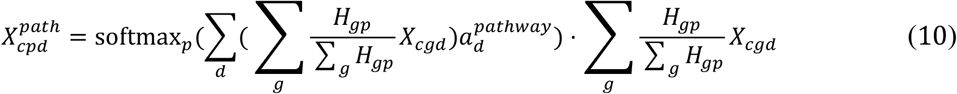

where *H*_*gp*_ is the gene–pathway incidence matrix encoding known gene membership in curated biological pathways, and 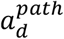 is a learnable pathway-level attention vector. This step aggregates gene embeddings into pathway representations for each cell, allowing the model to learn which pathways are most informative for characterizing transcriptional state. Pathway attention weights are defined as:

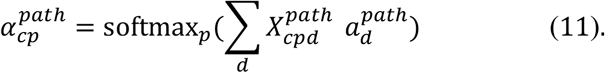

#### 2. Gene attention (pathway → gene message passing)

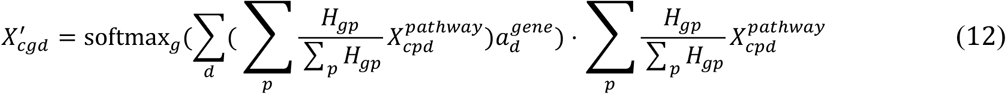

where 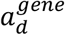 is a learnable gene-level attention vector. This step redistributes pathway-contextual information back to genes, enabling gene embeddings to incorporate signals from functionally related pathways. Biologically, this allows the model to emphasize genes that participate in coordinated pathway activity patterns associated with T cell activation, exhaustion, or cytotoxic response states. Gene attention weights are defined as:

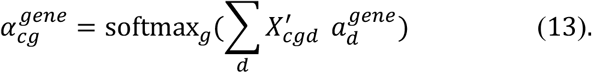

Residual connections and layer normalization were subsequently applied:

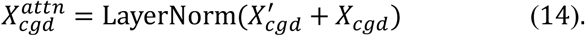

A feed-forward network (FFN) was then applied at each gene:

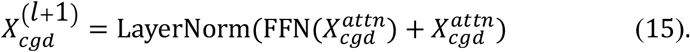

After six stacked hypergraph attention blocks, the encoder outputs the final gene-level cell embeddings:

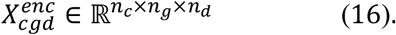

### Cell-level pooling

Gene-level embeddings were aggregated into cell-level embeddings using an expression-aware attention pooling mechanism. Let *E*_*cg*_ denote the normalized expression of gene *g* in cell *c*, and let 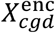 denote the corresponding encoded gene embedding. The pooled cell embedding is defined as:

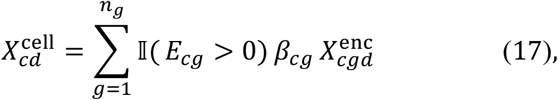

where I(⋅) is the indicator function and *β*_*cg*_ are learnable attention coefficients normalized across expressed genes within each cell.

The indicator function ensures that genes with zero observed expression do not contribute to the pooled cell representation. Consequently, attention normalization and feature aggregation are performed only across detected genes, naturally respecting the sparsity structure of scRNA-seq data. This expression-aware pooling strategy enables the model to emphasize transcriptionally active gene programs relevant to each cell state (such as activation, memory, or exhaustion programs in T and CAR T cells) while minimizing noise introduced by unexpressed genes.

### Training objectives

Module I was trained under a multi-task objective that jointly optimizes cell-type classification, gene expression reconstruction, and batch-adversarial domain adaptation.

#### Cell-type classification loss

A fully connected classifier predicts cell-type labels from cell embeddings. The loss is defined as cross-entropy between predicted and true cell-type labels:

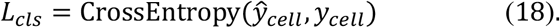

#### Gene expression reconstruction loss

A decoder reconstructs normalized gene expression from encoded gene-level embeddings. The reconstruction loss is defined as mean squared error:

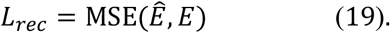

#### Batch-adversarial loss

A batch discriminator predicts dataset-of-origin labels from cell embeddings. The discriminator is optimized using cross-entropy:

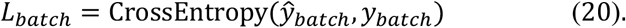

The total training objective for Module I is defined as:

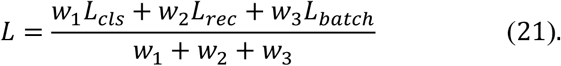

where *w*_1_, *w*_2_, *w*_3_ are loss weights. L2 regularization was applied to all trainable parameters.

### Batch effect correction via gradient reversal

To reduce dataset-specific technical variation while preserving biological signal, we adopted an adversarial domain adaptation strategy implemented through a gradient reversal layer ^52^. The batch discriminator was trained to correctly classify batch labels from cell embeddings, whereas the encoder was simultaneously optimized to prevent accurate batch discrimination. During backpropagation, gradients from the batch classification loss were multiplied by a negative constant −*γ* before propagating into the encoder. This adversarial optimization encourages batch-invariant representations while retaining biologically meaningful structure relevant to downstream prediction tasks.

Formally, discriminator parameters were optimized to minimize *L*_*batch*_, while encoder parameters are optimized to minimize *L*_*cls*_ + *L*_*rec*_ while maximize *L*_*batch*_ . The gradient reversal layer provides a stable implementation of this min-max objective.

### Training strategy and parameter settings of Module I

Model parameters were initialized using Xavier initialization and optimized using the Adam optimizer. The initial learning rate was set to 0.00001, with weight decay 0.0001. Training was performed using mini-batches of 64 cells. Dropout with rate 0.2 was applied in feed-forward layers to reduce overfitting. The hidden embedding dimension *n*_*d*_ was set to 128. Six stacked hierarchical hypergraph-attention blocks were used in the encoder.

Loss weights were set to *w*_1_ = 1, *w*_2_ = 1, and *w*_3_ = 0.35 for cell classification, reconstruction, and batch-adversarial objectives, respectively. The gradient reversal coefficient *γ* was set to 1. Models were trained for 10 epochs, and the checkpoint with the lowest validation loss was selected for downstream evaluation.

To adaptively adjust optimization dynamics during training, we used torch.optim.lr_scheduler.ReduceLROnPlateau. The learning rate reduction factor was set to 0.1, and the scheduler patience was set to 2 epochs without validation improvement. All models were implemented in PyTorch and trained on NVIDIA GPUs.

## Module II: Patient-level response prediction

Module II predicts patient-level CAR T therapy response using fixed cell embeddings learned from Module I. After training Module I, we obtained a cell embedding tensor 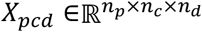, where *p, c*, and *d* index patients, cells, and embedding dimensions, respectively.

Parameters from Module I remained frozen during Module II training.

A cell-to-patient attention layer was used to aggregate cell information for each patient. Attention weights 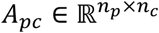 were computed as

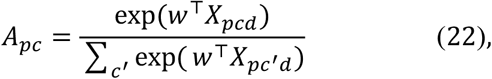

where 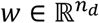 is a learnable attention vector and *X*_*pcd*_ denotes the embedding of cell *c* from patient *p*. The patient embedding matrix 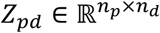 was then obtained by weighted aggregation:

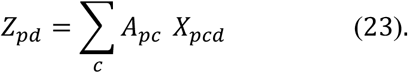

The resulting patient embeddings were passed through a multilayer perceptron classifier to generate predicted response probabilities:

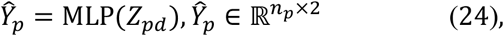

To account for class imbalance between responders and non-responders, Module II was trained using a weighted cross-entropy loss:

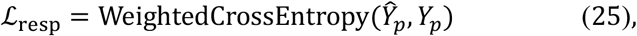

where loss weights for non-responding and responding classes were set to 3:1, reflecting the empirical class imbalance in the training cohort. Biologically, the cell-to-patient attention weights quantify the relative contribution of distinct CAR T cell states to patient-level therapeutic outcome.

### Implementation and training details of Module II

Module II was implemented in PyTorch and trained independently using fixed cell representations derived from each upstream model, including gANCHOR, Geneformer, scGPT, scMulan, scFoundation, GenePT, and raw gene expression profiles as an ablation baseline. To ensure fair comparison, all inputs were evaluated using the same downstream architecture and training protocol. To standardize patient-level input size, samples containing more than 2,000 cells were randomly subsampled to 2,000 cells for response prediction. The same sampled cells were used across all compared models to ensure fair and consistent benchmarking. We performed a grid search over optimization hyperparameters, including initial learning rate {1 × 10^−5^, 5 × 10^−5^, 1 × 10^−4^, 5 × 10^−4^, 1 × 10^−3^, 5 × 10^−3^, 1 × 10^−2^} and weight decay {10^−2^, 10^−3^, 10^−4^} . For each hyperparameter configuration, four independent runs with different random seeds were performed to account for stochastic optimization variability. Model selection was based on the average binary F1 score on the validation set across the four runs. The optimal hyperparameter configuration for each representation type was subsequently valuated on the held-out test cohort. Final reported performance metrics correspond to the distribution of test F1 scores across the four repeated runs.

The model was optimized using the Adam optimizer with early stopping based on validation loss to mitigate overfitting. All architectural components, including the attention-based aggregation layer and multilayer perceptron classifier, were kept identical across all experimental settings to ensure that observed performance differences reflect representation quality rather than downstream architectural variation.

### Model interpretability via hierarchical attention

gANCHOR provides interpretability across multiple biological resolutions through its hierarchical attention architecture. Gene-to-cell attention weights quantify the contribution of individual genes to cell embeddings, enabling identification of transcriptional programs associated with distinct cellular states. Cell-to-patient attention weights quantify the contribution of specific cellular populations to patient-level predictions. These hierarchical attention mechanisms enable systematic identification of biologically meaningful gene and cell signatures associated with therapeutic response.

## Code Availability

gANCHOR and all script files used in the analysis in this manuscript can be downloaded from GitHub repository at https://github.com/luoyuanlab/cart.

## Data Availability

All datasets used in this study are publicly available. T cell datasets were obtained from the CZ CELLxGENE database ^49^, including AIDA 2025 ^42^, Stephenson et al. 2021 ^43^, Yoshida et al. 2022 ^44^, Jin et al. 2021 ^45^, Rodríguez-Ubreva et al. 2022 ^46^, Chan Zuckerberg 2020 ^47^, and van der Wijst et al. 2021 ^48^, and are accessible via the following collection links: https://cellxgene.cziscience.com/collections/ced320a1-29f3-47c1-a735-513c7084d508, https://cellxgene.cziscience.com/collections/ddfad306-714d-4cc0-9985-d9072820c530, https://cellxgene.cziscience.com/collections/03f821b4-87be-4ff4-b65a-b5fc00061da7, https://cellxgene.cziscience.com/collections/b9fc3d70-5a72-4479-a046-c2cc1ab19efc, https://cellxgene.cziscience.com/collections/bf325905-5e8e-42e3-933d-9a9053e9af80, https://cellxgene.cziscience.com/collections/eb735cc9-d0a7-48fa-b255-db726bf365af, and https://cellxgene.cziscience.com/collections/7d7cabfd-1d1f-40af-96b7-26a0825a306d. CAR T cell datasets were obtained from the Gene Expression Omnibus (GEO) under accession numbers GSE151511 (Deng et al. 2020 ^17^), GSE197268 (Haradhvala et al. 2022 ^18^), GSE197215 (Bai et al. 2022 ^29^), and GSE262072 (Bai et al. 2024 ^30^). The dataset from Wilson et al. 2022 ^19^ is available from Dryad at https://datadryad.org/dataset/doi:10.5061/dryad.1rn8pk0x4.

