## Supplementary Information for "CAR T cell foundation model predicts immunotherapy response"

### Supplementary Text

#### T cell and CAR T cell single cell data collection and initial filtering

scRNA-seq datasets of T cells and CAR T cells were collected from a total of twelve published studies spanning multiple disease contexts. To support both representation learning and downstream clinical modeling, we curated two complementary data sources: (i) CAR T-cell datasets with patient-level clinical outcomes and pre-infusion/infusion-product profiles, and (ii) large-scale reference T-cell atlases capturing diverse functional states. Several CAR T source studies also collected longitudinal post-infusion samples; however, the CAR T-cell profiles used for Module I pretraining and Module II patient-level response prediction were derived from pre-infusion or infusion-product material collected before clinical outcome assessment, supporting analysis of transcriptional states associated with subsequent therapeutic response, persistence, and relapse [1-5]. In total, 161 patients across five CAR T studies were included for downstream patient-level analyses after applying study-specific filtering criteria. We first describe the CAR T-cell datasets used for patient-level modeling and evaluation, followed by the reference T-cell datasets used to support foundation model pretraining and to capture broad T-cell state diversity.

#### CAR T-cell datasets

Deng et al. 2020 [1] characteristics of anti-CD19 CAR T cell infusion products. CAR T-cell data were obtained from a study characterizing the infusion products of patients with large B-cell lymphoma (LBCL). The dataset comprises single-cell transcriptomic profiles of remnant cells from axicabtagene ciloleucel (axi-cel) infusion products administered to 24 patients with relapsed or refractory LBCL as standard-of-care therapy. All 24 patients were included in downstream analyses. Single-cell RNA sequencing was performed using the 10x

Genomics 5’ chemistry platform. This dataset provides reference profiles connecting specific CAR T-cell transcriptional states to clinical efficacy and toxicity outcomes.

Wilson et al. 2022 [2] CAR T-cell data were obtained from a study investigating transcriptional trajectories in pediatric patients with B-cell acute lymphoblastic leukemia (B-ALL). The source study profiled CD19-CAR T cells from pre-infusion good manufacturing practice (GMP) products and post-infusion peripheral blood and bone marrow samples, enabling examination of CAR T-cell states associated with persistence and clinical efficacy. For the response prediction analyses in the current study, only pre-infusion GMP product profiles with available response annotations were retained. Among the 16 patients reported, two patients lacking pre-infusion scRNA-seq data and two patients without response records were excluded, resulting in 12 patients included in the response prediction task. Because the publicly available data for this study were provided as normalized expression values, we first recovered raw count estimates to ensure compatibility with our uniform preprocessing pipeline.

Haradhvala et al. 2022 [3] CAR-T data were obtained from a study analyzing cellular dynamics in patients with refractory large B-cell lymphoma (LBCL) treated with CD19-targeted CAR T-cell therapy. The dataset comprises single-cell transcriptomic profiles from 105 samples from 32 patients, including infusion products and peripheral blood mononuclear cells (PBMCs) collected at pretreatment and post-treatment timepoints. For the current response prediction analyses, we retained infusion-product CAR T-cell profiles collected prior to clinical outcome assessment and did not use post-treatment PBMC samples as prediction inputs. One patient without scRNA-seq data was excluded, resulting in 31 patients included in the response prediction task.

Bai et al. 2022 [4] described a CAR T study analyzes the antigen-specific landscape of infusion products in patients with acute lymphoblastic leukemia (ALL). The dataset comprises 101,326 single-cell transcriptomes derived from pre-infusion CAR T-cell infusion products from 12 patients across basal and stimulation conditions. After study-specific filtering, all 12 patients were retained for downstream analyses, enabling a direct comparison between patients who experienced CD19-positive relapse and those achieving durable remission.

Bai et al. 2024 [5] CAR T atlas designed to uncover the determinants of ultra-long-term remission in pediatric B-ALL. This dataset comprises 695,819 scRNA-seq profiles from 82 patients, capturing pre-infusion CAR T cells under both basal conditions and following CAR-specific stimulation. After study-specific filtering, all 82 patients were retained for downstream analyses.

To complement the relatively limited number of CAR T patient samples and to improve the robustness and generalizability of learned representations, we further incorporated large-scale reference T-cell datasets spanning multiple physiological and disease conditions. These datasets provide extensive coverage of T-cell functional heterogeneity, including naive, memory, cytotoxic, and regulatory T-cell states, thereby serving as a foundation for learning biologically meaningful cell representations.

**Reference T-cell datasets**

T cell datasets were obtained from large-scale atlases and disease-focused cohorts and were subsetted to retain annotated T cell populations across diverse functional subtypes [6-12].

AIDA 2025 [6] provides T cell data representing healthy baseline conditions, obtained from the Asian Immune Diversity Atlas and accessed via the CZ CELLxGENE database [13]. The source dataset comprises peripheral blood mononuclear cells (PBMCs) from 619 healthy donors spanning seven population groups across five Asian countries, profiled using 10x Genomics 5' scRNA-seq. For the current study, we specifically subsetted the data to retain only T cell populations, filtering for the following subtypes: CD8-positive alpha-beta memory T cell, regulatory T cell, central memory CD4-positive alpha-beta T cell, naive thymus-derived CD4-positive alpha-beta T cell, T cell, naive thymus-derived CD8-positive alpha-beta T cell, mucosal invariant T cell, gamma-delta T cell, CD8-positive alpha-beta cytotoxic T cell, CD4-positive alpha-beta cytotoxic T cell, effector memory CD4-positive alpha-beta T cell, CD4-positive alpha-beta T cell, CD8-positive alpha-beta T cell, and double negative T regulatory cell.

Stephenson et al. 2021 [7] profiles T cell responses in COVID-19 through a single-cell multi-omics study, with data accessed via the CZ CELLxGENE database. The original dataset comprises over 780,000 peripheral blood mononuclear cells (PBMCs) from a cross-sectional cohort of 130 patients with varying disease severities. For this study, we subsetted the data to specifically retain T cell populations, filtering for the following subtypes: regulatory T cell, T-helper 22 cell, naive thymus-derived CD8-positive alpha-beta T cell, naive thymus-derived CD4-positive alpha-beta T cell, mature NK T cell, effector memory CD8-positive alpha-beta T cell, central memory CD4-positive alpha-beta T cell, mucosal invariant T cell, T follicular helper cell, gamma-delta T cell, effector memory CD4-positive alpha-beta T cell, effector CD8-positive alpha-beta T cell, T-helper 1 cell, T-helper 2 cell, T-helper 17 cell, CD8-positive alpha-beta T cell, and CD4-positive alpha-beta T cell.

Yoshida et al. 2022 [8] characterizes T cell data across pediatric and adult individuals using a single-cell multi-omic framework, with data accessed via the CZ CELLxGENE database. The original dataset includes matched nasal, tracheal, bronchial, and blood samples from 93 individuals, comprising pediatric and adult COVID-19 patients as well as healthy controls. For this study, we subsetted the data to specifically retain T cell populations, filtering for the following subtypes: naive thymus-derived CD8-positive alpha-beta T cell, naive thymus-derived CD4-positive alpha-beta T cell, CD4-positive alpha-beta cytotoxic T cell, mature NK T cell, gamma-delta T cell, CD4-positive helper T cell, CD8-positive alpha-beta cytotoxic T cell, effector memory CD8-positive alpha-beta T cell, regulatory T cell, mucosal invariant T cell, effector memory CD8-positive alpha-beta T cell terminally differentiated, and central memory CD8-positive alpha-beta T cell.

Jin et al. 2021 [9] presents a harmonized single-cell transcriptomic analysis of host responses to SARS-CoV-2 infection, from which T cell data were obtained via the CZ CELLxGENE database. The original dataset integrates data from blood, bronchoalveolar lavage, and tissue samples across COVID-19 patients and control conditions. For this study, we subsetted the data to specifically retain T cell populations, filtering for the following subtypes: central memory CD4-positive alpha-beta T cell, effector memory CD8-positive alpha-beta T cell, CD4-positive alpha-beta cytotoxic T cell, activated CD8-positive alpha-beta T cell human, regulatory T cell, naive thymus-derived CD4-positive alpha-beta T cell, naive thymus-derived CD8-positive alpha-beta T cell, activated CD4-positive alpha-beta T cell human, gamma-delta T cell, mucosal invariant T cell, and double negative thymocyte.

Rodríguez-Ubreva et al. 2022 [10] investigates T cell alterations in common variable immunodeficiency (CVID) using a single-cell multi-omics approach, with data accessed via the CZ CELLxGENE database. The original dataset comprises single-cell profiles from a CVID-discordant monozygotic twin pair, validated in a larger cohort of CVID patients and healthy donors. For this study, we subsetted the data to specifically retain T cell populations, filtering for the following subtypes: naive thymus-derived CD4-positive alpha-beta T cell, naive thymus-derived CD8-positive alpha-beta T cell, type I NK T cell, gamma-delta T cell, regulatory T cell, CD8-positive alpha-beta memory T cell, CD4-positive alpha-beta memory T cell, activated CD4-positive alpha-beta T cell, activated CD8-positive alpha-beta T cell, mucosal invariant T cell, effector memory CD8-positive alpha-beta T cell terminally differentiated, and effector memory CD8-positive alpha-beta T cell.

Chan Zuckerberg 2020 [11] provides an international multi-tissue single-cell resource of COVID-19 patients, from which T cell data were obtained via the CZ CELLxGENE database. The original dataset includes peripheral blood and nasal swab samples from patients with COVID-19 who had pre-existing immunological conditions, including autoimmune diseases (rheumatoid arthritis, psoriasis, Sjogren syndrome, eosinophilic granulomatosis with polyangiitis, multiple sclerosis) and common variable immunodeficiency (CVID). For this study, we subsetted the data to specifically retain T cell populations, filtering for the following subtypes: CD8-positive alpha-beta memory T cell, CD4-positive alpha-beta memory T cell, mucosal invariant T cell, mature NK T cell, gamma-delta T cell, naive thymus-derived CD4-positive alpha-beta T cell, naive thymus-derived CD8-positive alpha-beta T cell, regulatory T cell, and T cell.

van der Wijst et al. 2021 [12] examines the role of type I interferon autoantibodies in COVID-19 using longitudinal CITE-seq data, from which T cell profiles were obtained via the CZ CELLxGENE database. The original dataset comprises longitudinal single-cell epitope and transcriptome sequencing (CITE-seq) profiles of peripheral blood mononuclear cells (PBMCs) from 54 patients with varying degrees of COVID-19 severity and 26 non-COVID-19 controls. For this study, we subsetted the data to specifically retain T cell populations, filtering for the following subtypes: CD4-positive alpha-beta memory T cell, CD4-positive alpha-beta T cell, regulatory T cell, mucosal invariant T cell, gamma-delta T cell, CD8-positive alpha-beta memory T cell, CD8-positive alpha-beta T cell, and mature NK T cell.

**Benchmark single cell foundation models**

To systematically evaluate model performance, we benchmarked gANCHOR against five representative single-cell foundation models spanning gene-level and cell-level representation learning paradigms. All methods were applied using publicly available pretrained models with default parameterizations to ensure consistency and reproducibility. For each method, embeddings were extracted and directly used as input to the same downstream patient-level prediction framework.

Geneformer [14] is a transformer-based foundation model pretrained on large-scale single-cell transcriptomic data to learn contextualized gene representations through masked language modeling. It encodes gene expression profiles as ranked gene tokens and captures gene-gene dependencies using self-attention mechanisms. In this study, we ran Geneformer using its official implementation. We used the publicly available pretrained Geneformer model with default parameterization to extract cell embeddings for CAR T cells, which were subsequently used as input to the downstream response prediction module. The resulting cell embedding dimension is 256.

scGPT [15] is a generative pre-trained transformer model designed for single-cell multi-omics analysis. It models gene expression profiles using a transformer architecture with autoregressive and masked modeling objectives, enabling the learning of scalable and transferable cell representations. We applied the pretrained scGPT model with default settings to obtain cell embeddings for CAR T cells without additional fine-tuning. These embeddings were directly used for downstream patient-level prediction. The resulting cell embedding dimension is 512.

GenePT [16] is a gene embedding framework that leverages large language models to encode gene-level biological knowledge from textual corpora. Unlike cell-level foundation models, GenePT provides fixed gene embeddings rather than cell embeddings. In this study, we directly extracted pretrained gene embeddings and used them to represent single-cell profiles by mapping gene expression values onto the corresponding embedding space. The resulting representations were used as input features for downstream analysis. The gene embedding dimension is 1,536.

scMulan [17] is a multitask generative pre-trained model for single-cell analysis that integrates multiple learning objectives, including gene expression modeling and cell-type prediction, to learn robust cell representations. The model employs a transformer-based architecture to capture complex gene dependencies and supports cross-dataset generalization. We used the publicly available pretrained scMulan model with default parameter settings to extract CAR T-cell embeddings for downstream response prediction. The resulting cell embedding dimension is 1,120.

scFoundation [18] is a large-scale foundation model trained on extensive single-cell transcriptomic datasets to learn generalizable cell representations. It utilizes a transformer-based architecture optimized for scalability and transfer learning across diverse biological contexts. In this study, we applied the pretrained scFoundation model with default configurations to generate cell embeddings for CAR T cells, which were then used in the unified downstream prediction framework. The resulting cell embedding dimension is 3,072.

### Supplementary Figures

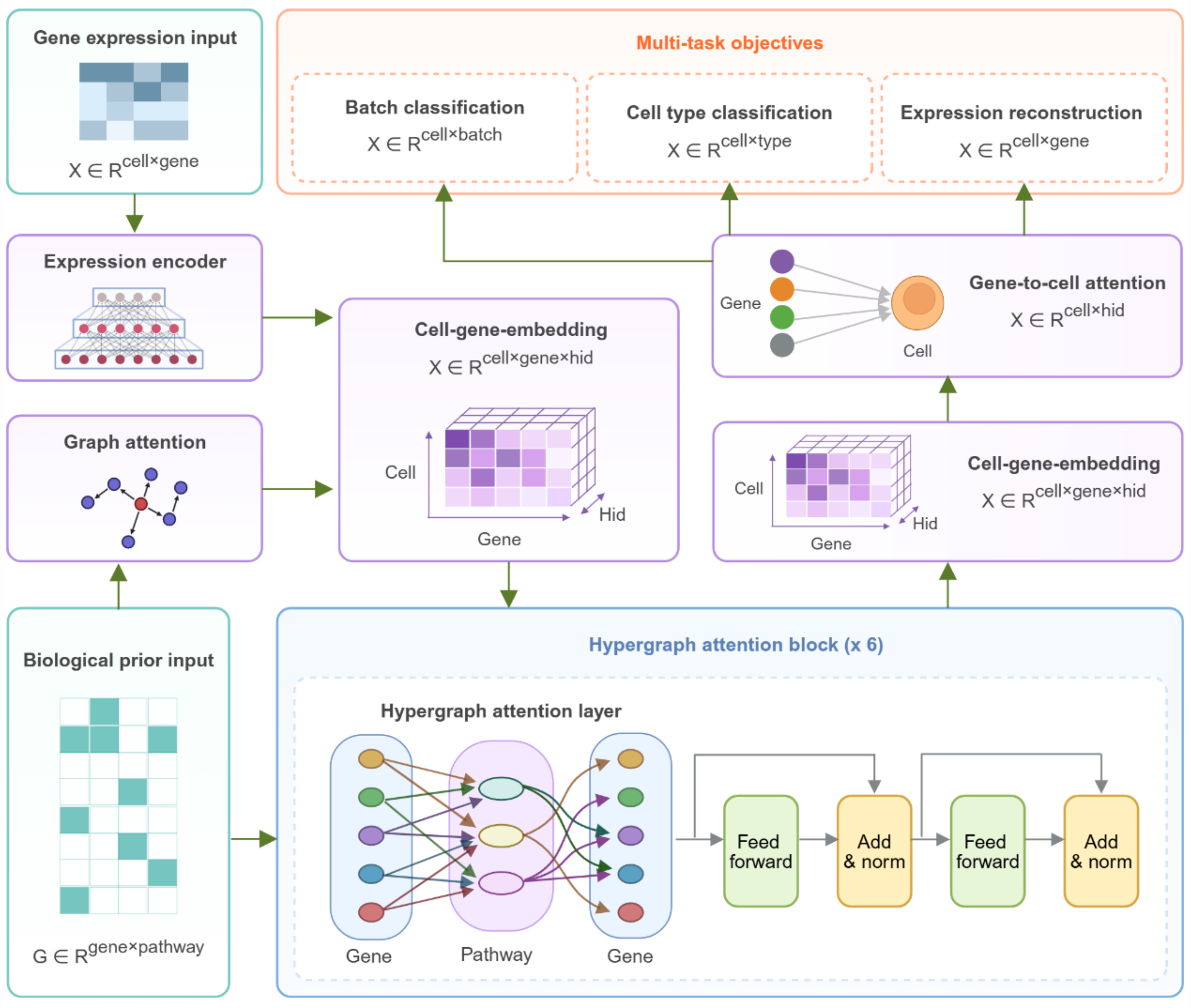


**Figure S1 Architecture of Module I in gANCHOR.** Schematic overview of Module I in gANCHOR, illustrating the flow of information from input gene expression to multi-task outputs. Single-cell gene expression profiles, together with the biological prior gene-pathway association matrix, are first processed by an expression encoder and graph attention layers to generate initial cell-gene embeddings, which are subsequently refined through stacked hypergraph attention blocks that perform gene-pathway-gene message passing to capture structured biological dependencies. The refined embeddings are then aggregated via a gene-to-cell attention mechanism to produce cell-level representations, which are used for multi-task learning objectives, including cell-type classification, batch classification (via adversarial learning), and gene expression reconstruction.

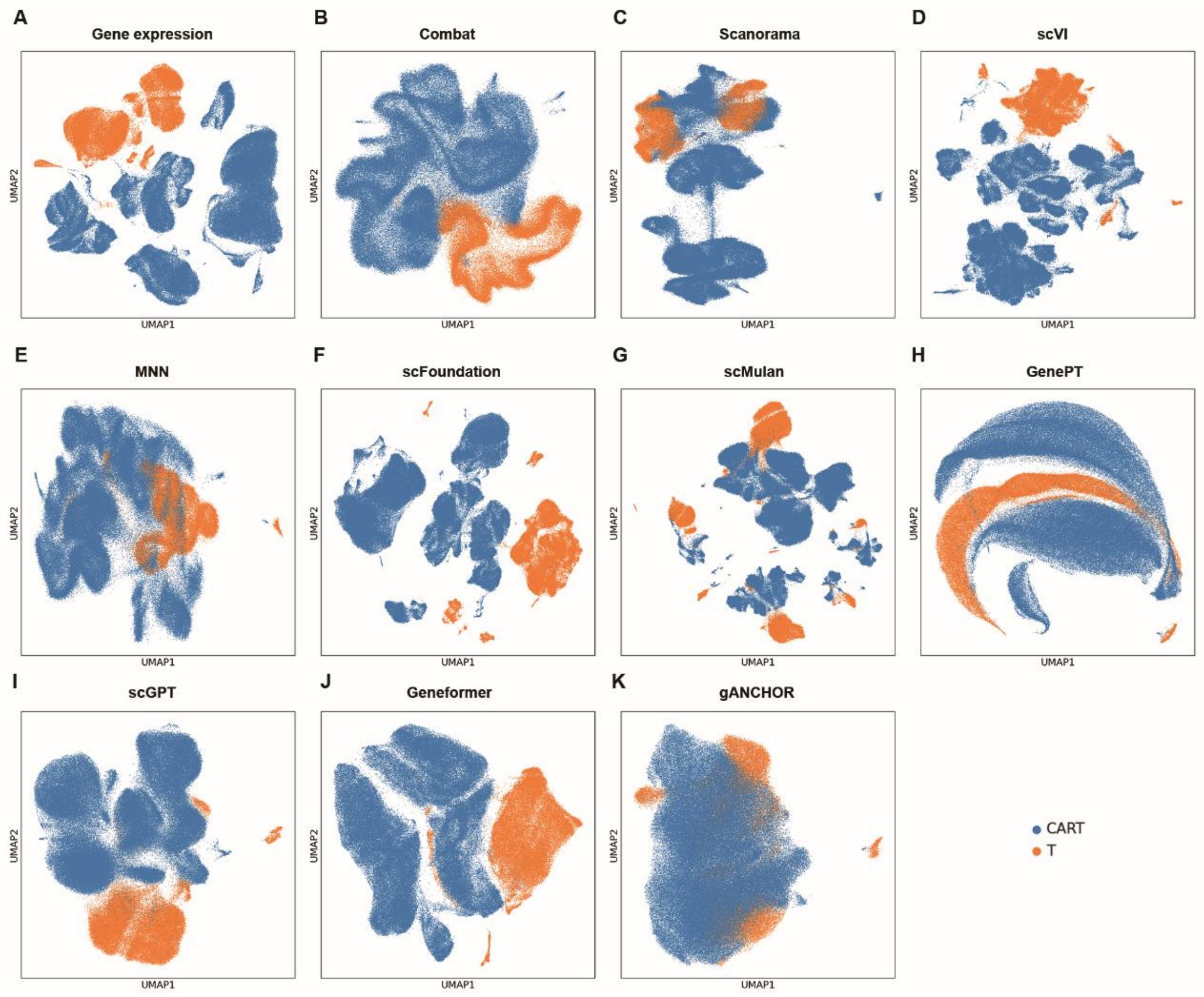


**Figure S2 UMAP visualization of cell embeddings.** UMAP projections of cell embeddings generated from raw gene expression (**A**), Combat (**B**), Scanorama (**C**), scVI (**D**), mutual nearest neighbor (MNN) (**E**), scFoundation (**F**), scMulan (**G**), GenePT (**H**), scGPT (**I**), Geneformer (**J**), and gANCHOR (**K**), with cells colored by dataset origin (CAR T vs. T cell datasets). Compared to other approaches, gANCHOR demonstrates improved integration between CAR T and T cell datasets while maintaining coherent global structure.

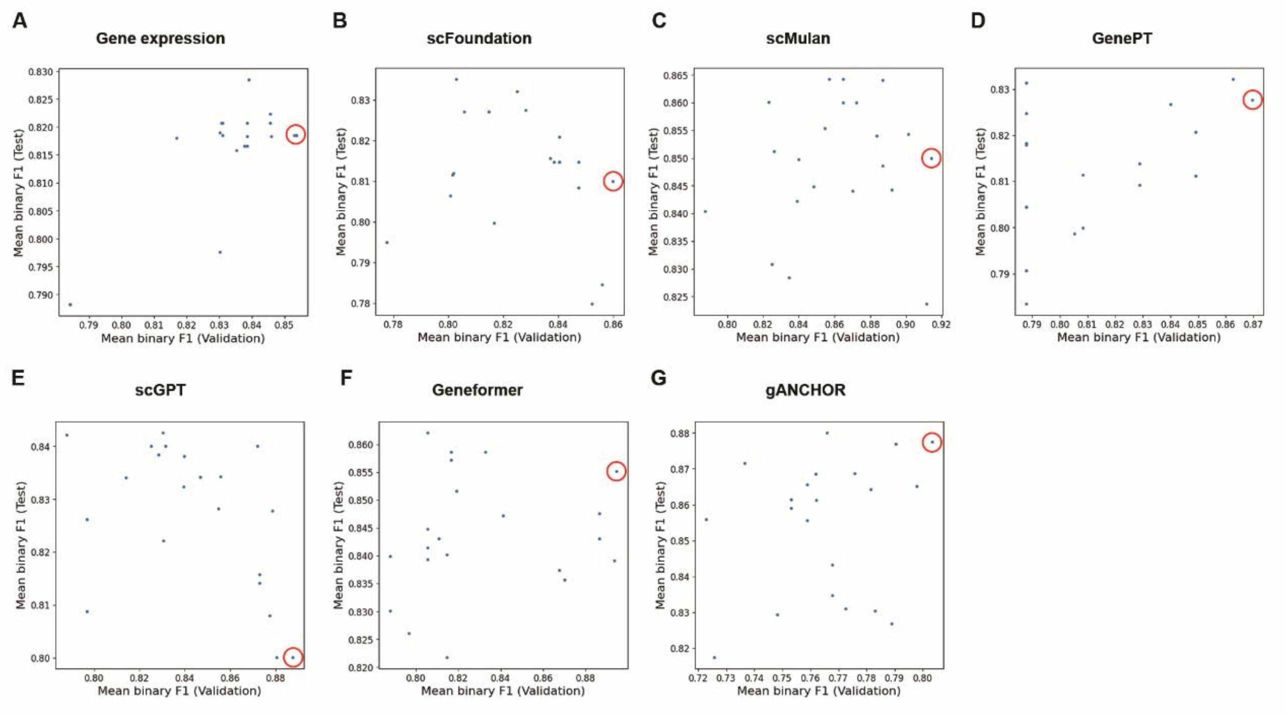


**Figure S3 Hyperparameter search and performance of benchmark models.** (**A-G**) Scatter plots showing the relationship between validation and test performance across hyperparameter configurations for each model, including raw gene expression (**A**), scFoundation (**B**), scMulan (**C**), GenePT (**D**), scGPT (**E**), Geneformer (**F**), and gANCHOR (**G**). Each point represents one hyperparameter configuration, with coordinates corresponding to the mean binary F1 score on the validation set (x-axis) and the test set (y-axis), averaged over four independent training runs with different random seeds. A grid search was performed over optimization hyperparameters, including initial learning rates $\{1\times10^{-5}, 5\times10^{-5}, 1\times10^{-4}, 5\times10^{-4}, 1\times10^{-3}, 5\times10^{-3}, 1\times10^{-2}\}$ and weight decay values $\{10^{-2}, 10^{-3}, 10^{-4}\}$, resulting in 21 total configurations per model. Red circles indicate the configuration with the highest validation F1 score, which was selected for final evaluation on the test set.

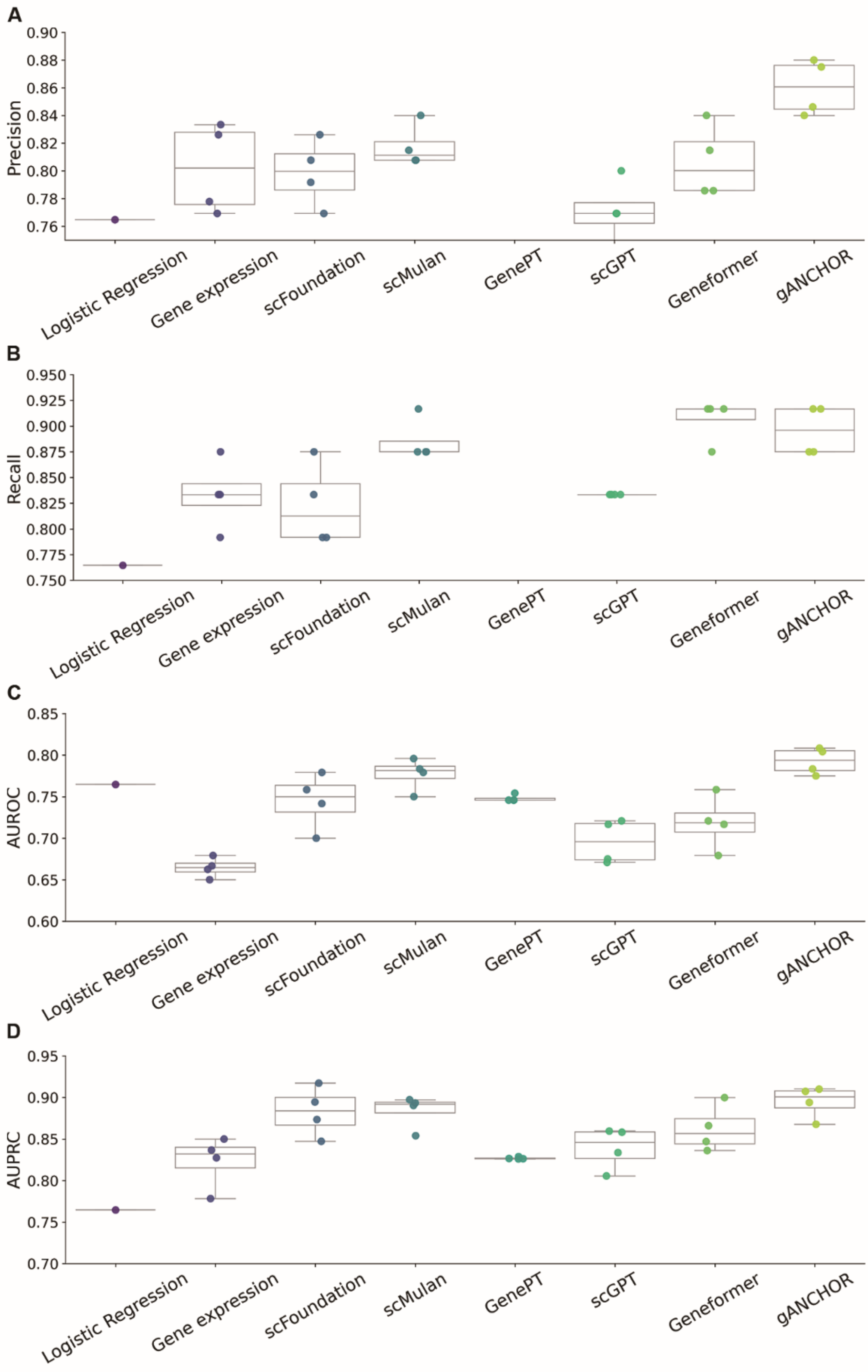


**Figure S4. Evaluation metrics for patient-level CAR T therapy response prediction.** Comparison of predictive performance across gANCHOR and benchmark models on the held-out test set using **(A)** precision, **(B)** recall, **(C)** area under the receiver operating characteristic curve (AUROC), and **(D)** area under the precision–recall curve (AUPRC). Each point represents the performance from an independent training run using a different random seed, and box plots summarize the distribution across four runs.

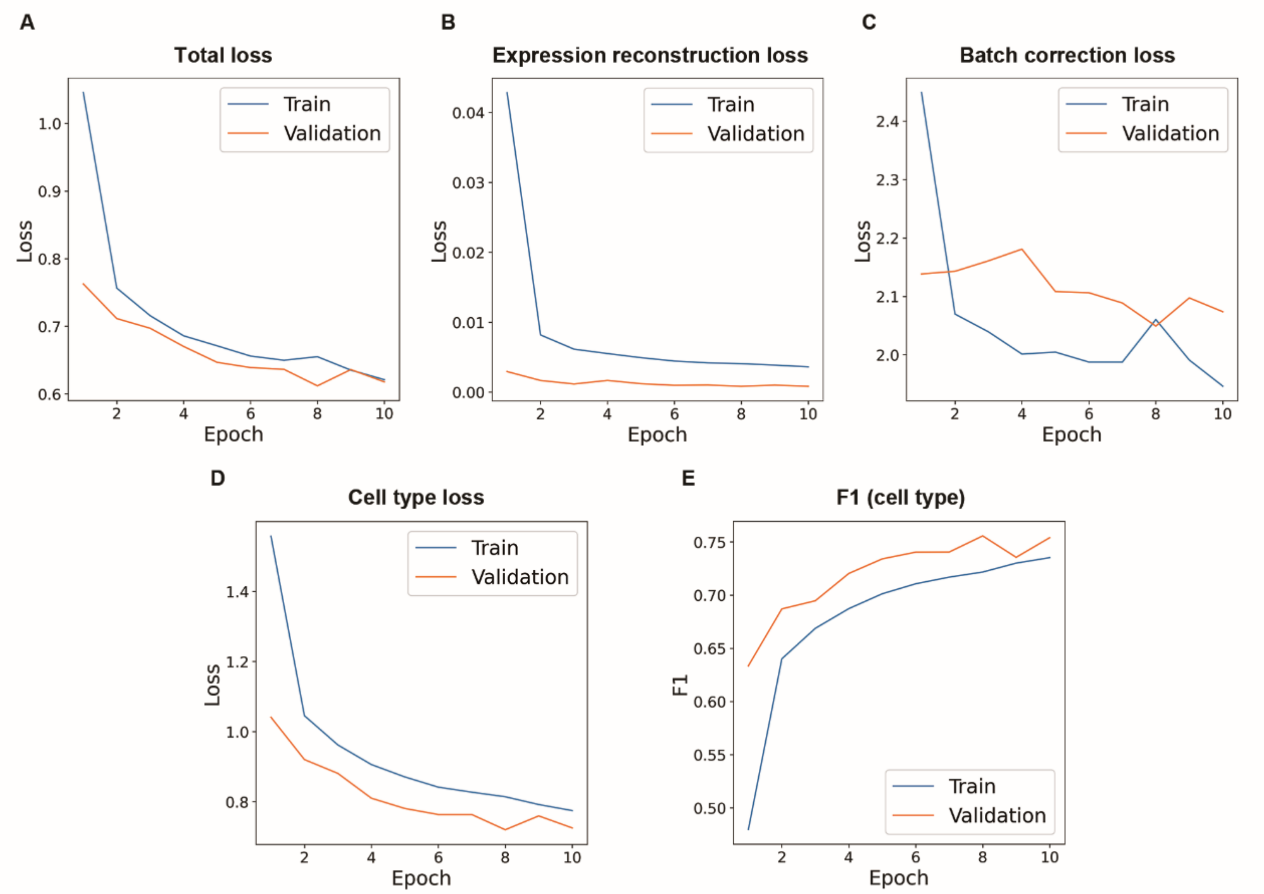


**Figure S5 Training dynamics of Module I.** (**A-D**) Training and validation loss curves over epochs for total loss (**A**), expression reconstruction loss (**B**), batch correction loss (**C**), and cell type classification loss (**D**). (**E**) Corresponding F1 score for cell type prediction on training and validation sets. The model was trained using a multi-task loss combining reconstruction, cell classification, and batch-adversarial components, with early stopping based on validation performance. These results demonstrate stable convergence and consistent improvement in cell type prediction performance during training.

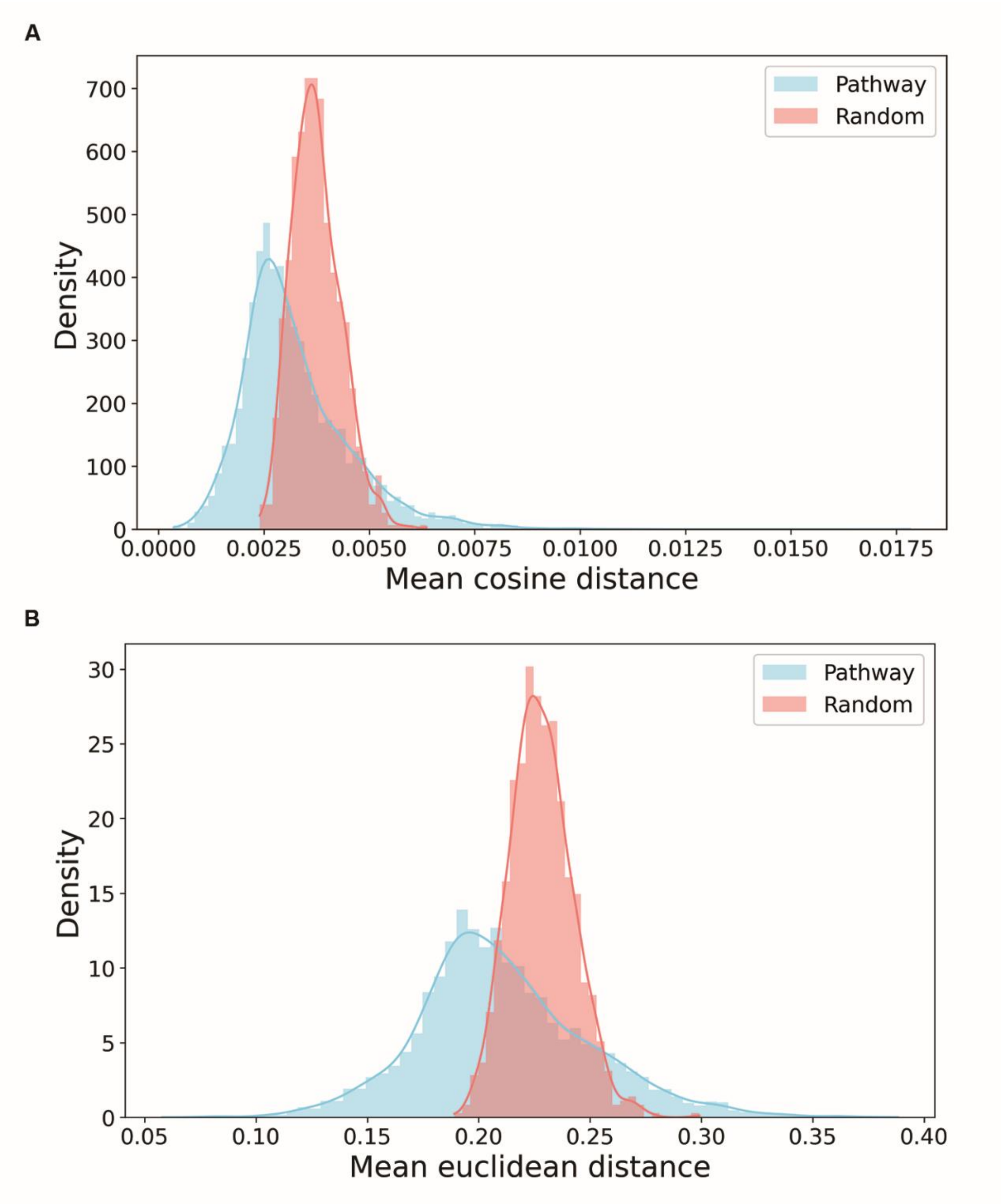


**Figure S6 Distribution of gene–gene distances within pathways and random gene sets.** (**A, B**) Distributions of mean pairwise cosine (**A**) and Euclidean (**B**) distances for genes within the same pathways (blue) compared to randomly sampled gene groups (red). Random groups were generated by sampling 1,000 gene sets matched in size to pathway gene sets. Genes within the same pathway consistently exhibit smaller pairwise distances than random gene groups.

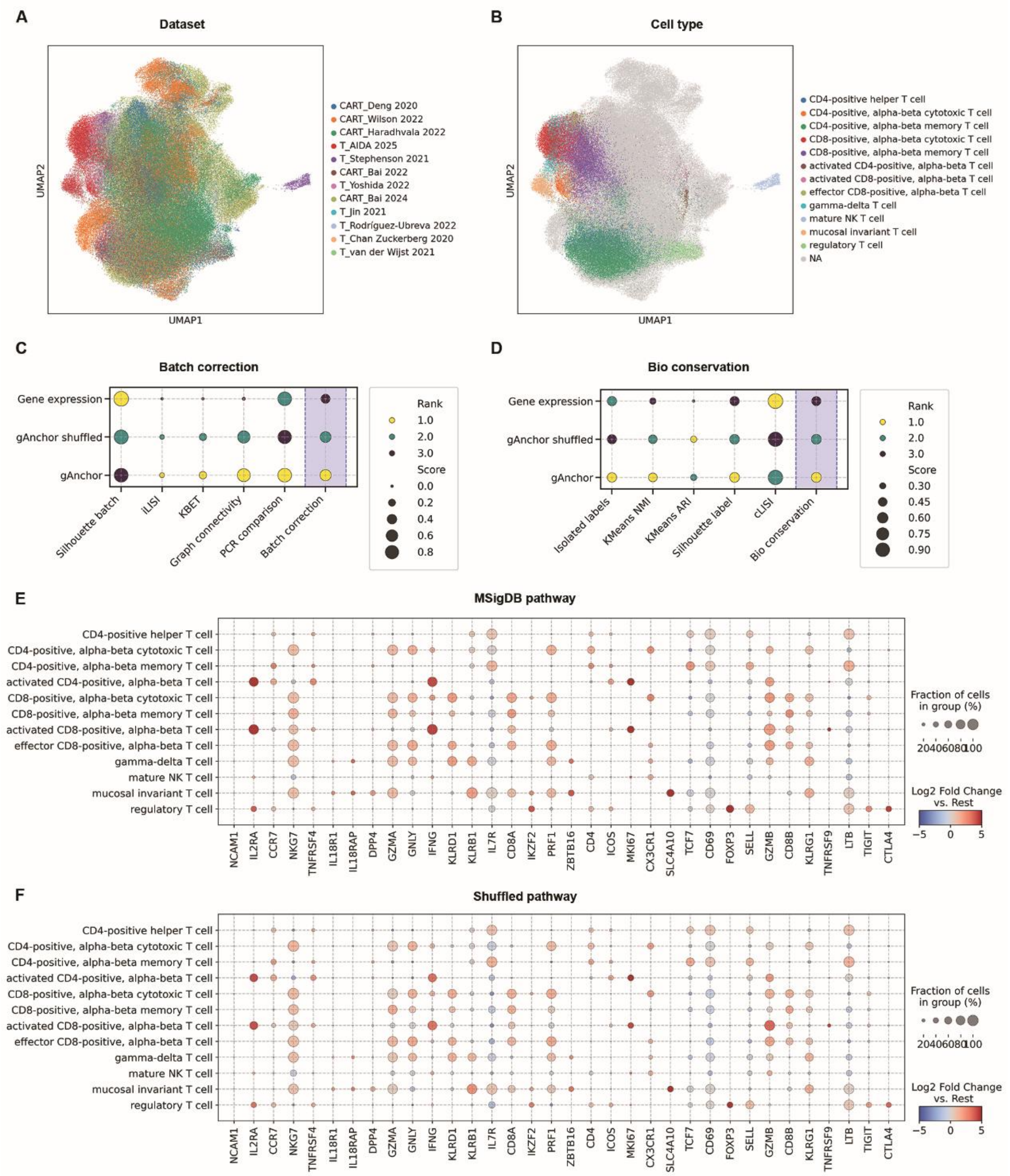


**Figure S7. Ablation analysis demonstrating the contribution of curated gene-pathway relationships to representation learning.** (**A**) UMAP visualization of cell embeddings generated using a shuffled gene-pathway graph, colored by dataset. (**B**) The same embedding colored by annotated T-cell subtype. (**C, D**) Comparison of batch-correction metrics (**C**) and biological conservation metrics (**D**) between gANCHOR trained with the curated MSigDB pathway graph and the shuffled pathway graph. (**E, F**) Dot plot of canonical T-cell marker genes ranked by differential attention-enriched genes using the curated MSigDB pathway graph (**E**) and shuffled pathway graph (**F**). Dot size indicates the fraction of cells with high attention scores, and color represents the log2 fold change in attention score relative to all remaining cells.

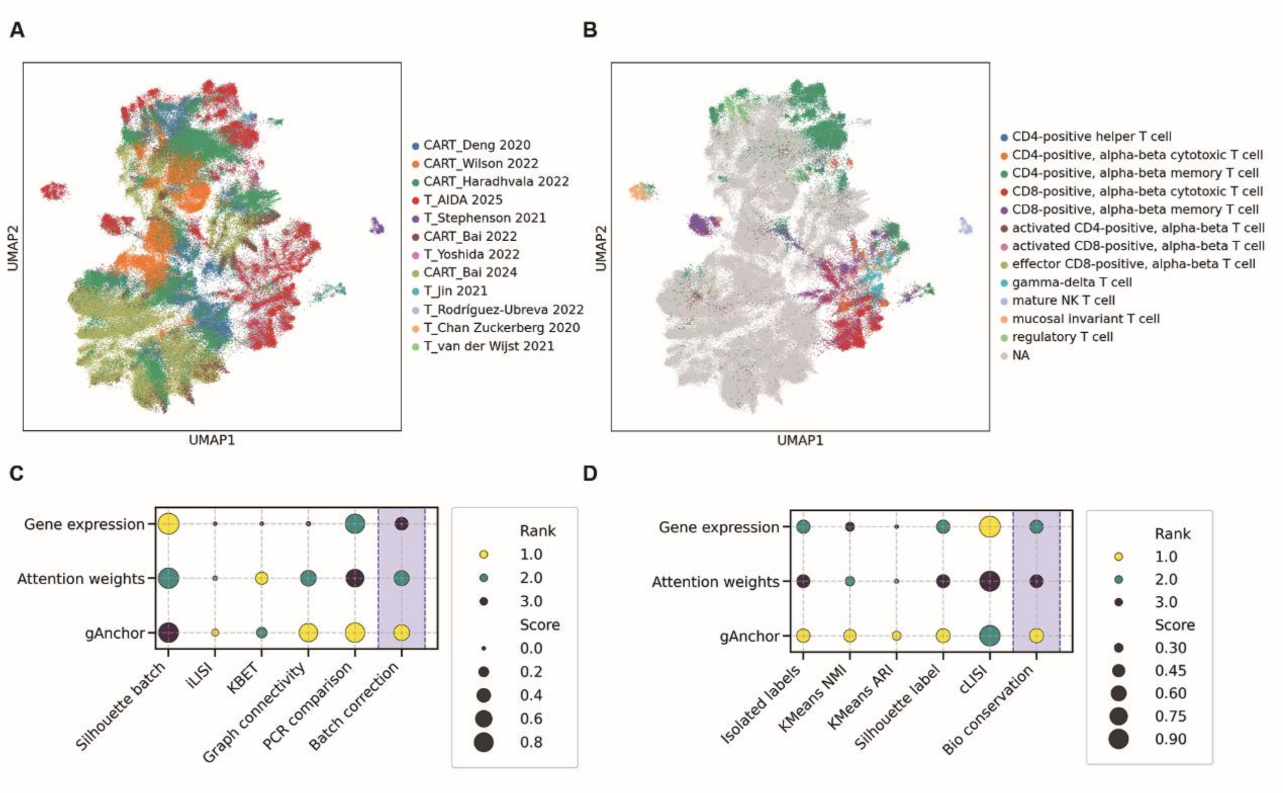


**Figure S8 Attention-derived representations preserve biologically structured cellular states.** (**A, B**) UMAP visualization of gene-to-cell attention-derived representations extracted from the attention module of gANCHOR. Cells are colored by dataset identity (**A**) and annotated T-cell subtype (**B**). Grey cells in (**B**) indicate CAR T cells without subtype annotations. (**C, D**) Quantitative comparison of normalized gene expression, attention-derived representations, and gANCHOR cell embeddings across batch correction (**C**) biological conservation (**D**) metrics. Dot size represents normalized metric score and color indicates rank among representations.

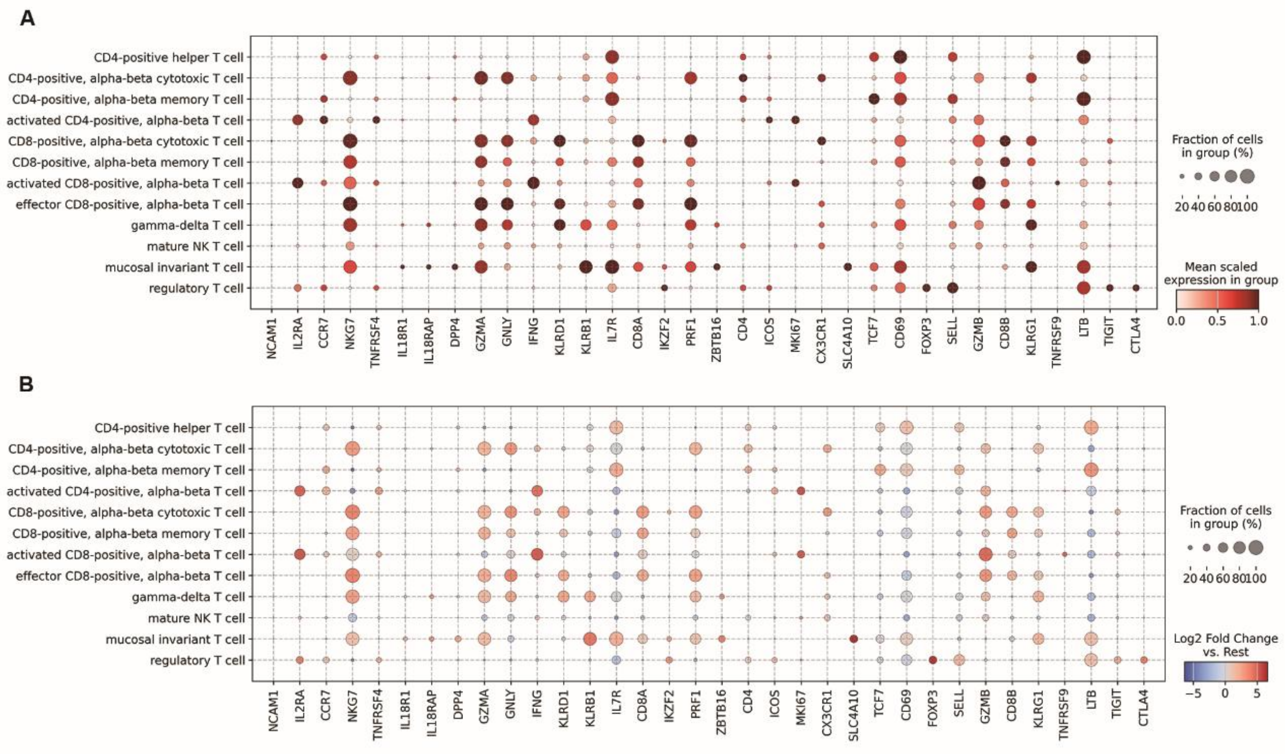


**Figure S9. Differential expression analysis identifies canonical marker genes across T-cell subtypes. (A)** Dot plot showing the expression patterns of representative marker genes identified by conventional differential expression analysis across annotated T-cell subtypes. Dot size indicates the percentage of cells expressing each gene, and color represents the mean scaled expression level within each cell type. **(B)** Corresponding differential expression statistics for the same marker genes. Dot size indicates the percentage of expressing cells, and color denotes the log2 fold change relative to all remaining T-cell populations (one-versus-rest comparison).
